# How Reliable Are Reconstructions of Complex Polyploid Histories? A Benchmark of Phylogenetic Inference Methods

**DOI:** 10.64898/2026.11.26.769415

**Authors:** Tom Ulanovski, Ronen Shtein, Jérôme Tubiana, Tal Pupko, Itay Mayrose

## Abstract

Polyploid phylogenetic inference methods are regularly applied to reconstruct allo- and autopolyploidization events in clades with complex evolutionary histories. To date, however, they have been evaluated mainly on small, simulated networks containing a handful of species and very few polyploidization events. Moreover, the extent to which different methods recover the same evolutionary history from the same data is largely unexplored. Here, we benchmark five existing methods, PADRE, AlloppNET, GRAMPA, MPAllopp and Polyphest, together with GRAMPA^Iter^, an automated extension of GRAMPA developed here that enables its application to histories involving more than a single polyploidization event. We evaluated their agreement on 12 empirical datasets and their accuracy on simulations based on 21 published phylogenetic networks encompassing dozens of species. Simulations varied incomplete lineage sorting, rates of gene duplication and loss, and extent of fractionation. To compare inferred networks, we introduce two measures that separately quantify disagreement in the descendant taxa and parental lineages associated with reticulation events. Only Polyphest, GRAMPA and GRAMPA^Iter^ consistently completed analyses within practical runtimes. On empirical data, the methods differed substantially in which species they inferred to be polyploid and in the number and placement of reticulations. Polyphest was the most accurate method in all 12 simulated configurations without fractionation. Under strong fractionation, however, its accuracy declined sharply, whereas GRAMPA^Iter^, which does not require ploidy, was considerably more robust. Our results reveal substantial uncertainty in reconstructing complex polyploid histories, with performance depending strongly on the evolutionary processes shaping the data.

## Main

Polyploidy, the presence of more than two complete sets of chromosomes in the genome of an organism, is a major evolutionary process in plants and certain other eukaryotic groups such as fish and insects (Otto and Whitton 2000; Mable et al. 2011; Van de Peer et al. 2017). Whole-genome duplications (WGDs) are widespread throughout plant evolution with most, if not all, plant lineages showing evidence of at least one WGD in their evolutionary history (Cui et al. 2006; Jiao et al. 2011) and an estimated 30–35% of extant flowering plant species are of recent polyploid origin (Wood et al. 2009; Halabi et al. 2023). Polyploidy can drive phenotypic changes and has been associated with increased adaptability and evolutionary innovation, with polyploid organisms demonstrating enhanced biological plasticity, stress resistance, and ability to colonize new ecological niches (Anatskaya and Vinogradov 2022). Understanding the evolutionary history of polyploid lineages therefore requires reconstructing the polyploidization events that gave rise to them.

Polyploidization is commonly classified into two main types. Autopolyploidy results from genome duplication within a single lineage, leading to nearly identical duplicated chromosome sets, whereas allopolyploidy arises through hybridization between distinct lineages, combining genomes with different evolutionary histories (Ramsey and Schemske 1998). Reconstructing the evolutionary history of polyploid lineages presents computational challenges due to processes including hybridization, genome doubling, and the subsequent gradual return towards a diploid state, a process termed diploidization, which can obscure evidence of ancient polyploidization events (Mandáková and Lysak 2018).

Allopolyploid evolution cannot be adequately represented by a conventional phylogenetic tree, in which each lineage has only one parent, whereas an allopolyploid lineage inherits genomes from two distinct parental lineages. Representing such evolutionary histories therefore requires a reticulate structure, such as a phylogenetic network, which accommodates contributions from multiple parental lineages (Fig. 1). A closely related representation, used by most of the methods considered here, is the multi-labeled tree (MUL-tree). In a MUL-tree, a species may appear more than once, with the number of tips reflecting the number of distinct subgenomes: a diploid is represented by one tip, a tetraploid by two, a hexaploid by three, and so on (Fig. 1b). MUL-trees and phylogenetic networks provide closely related representations of polyploid evolutionary histories. A MUL-tree explicitly represents the different homeologous genomes (i.e., subgenomes derived from different parental lineages) as separate copies of the same taxon, whereas a phylogenetic network represents their common polyploid lineage through reticulation (Figs. 1b, 1c).

**Figure 1.**
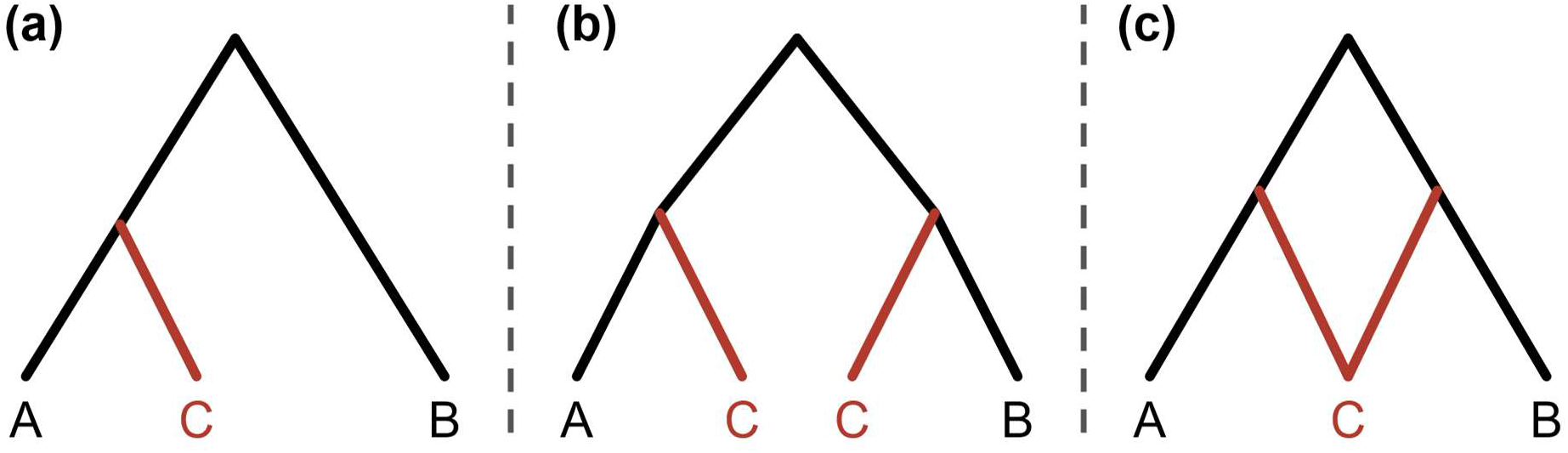
Alternative phylogenetic representations of allopolyploid evolution. In all panels, species C is an allopolyploid derived from parental lineages A and B. (a) A conventional species phylogeny is represented as a singly labeled tree, in which each species appears once. For a polyploid, this representation captures only one of its two parental histories, grouping C with either A or B. (b) A multi-labeled tree (MUL-tree) represents both subgenomes of C, with one copy descending from each parental lineage. As in a conventional tree, each node has an in-degree of one. (c) A phylogenetic network represents C with a single tip and the contribution of both parental lineages through a reticulation node with an in-degree of two, representing the hybridization event.

The inference of polyploid evolutionary histories from gene trees is particularly complicated by incomplete lineage sorting (ILS) and gene duplication and loss. ILS occurs when ancestral genetic variation persists across speciation events, such that the order in which gene lineages coalesce differs from the order in which the species diverged (Maddison 1997). It can therefore generate discordance between gene trees and the species history without altering the number of gene copies. Gene duplication and loss provide a second source of discordance and, unlike ILS, can alter gene copy number. Gene loss is particularly common during diploidization (Freeling 2009; Li et al. 2021), while small-scale duplications can introduce additional copies. When a loss in one subgenome is offset by a duplication in another, however, the total copy number may remain unchanged, making the resulting discordance difficult to distinguish from ILS.

General phylogenetic network methods, such as SNaQ (Solís-Lemus and Ané 2016) and several methods implemented in PhyloNet (Yu and Nakhleh 2015; Wen et al. 2018), were primarily developed to model hybridization among diploid lineages, without explicitly modeling genome duplication and the duplicated gene copies generated by polyploidization. These PhyloNet methods treat such copies as alleles of a single population, which are expected under the model to frequently coalesce within the polyploid lineage. Homeologs that trace to different parental lineages may therefore be represented as deep coalescences, increasing the inferred cost of the true polyploid history (Yan et al. 2022). SNaQ infers species networks from quartets of distinct species, in which each gene contributes equally regardless of how many copies a polyploid carries, so variation in copy number associated with genome duplication is not explicitly represented. It is further restricted to level-1 networks, in which each edge belongs to at most one reticulation cycle, whereas polyploid histories can violate this restriction, for example, when an allopolyploid itself contributes to a later polyploidization event. These differences in model scope motivated the development of specialized methods explicitly designed for polyploid phylogenetic reconstruction.

Despite their increasing use in empirical studies, polyploid-specific inference methods have received limited comparative evaluation across realistic evolutionary scenarios, with previous assessments generally restricted to relatively simple evolutionary histories involving few taxa and polyploidization events. Their performance in the presence of major sources of gene-tree discordance, and the extent to which different methods recover the same evolutionary history from the same data remain largely unexplored. Moreover, no standardized framework exists for comparing inferred polyploid networks. Existing comparison measures largely summarize overall topological dissimilarity (e.g., Yan et al. 2024), which does not reveal whether disagreement arises from which taxa are inferred to be polyploid, their parental lineages, or the presence of spurious or missing reticulation events.

Here, we systematically benchmark five existing methods representing different approaches to polyploid phylogenetic inference. We evaluate their accuracy using simulated data, where the true network is known, and their agreement across empirical datasets representing a diverse range of real-world polyploid histories. One of these methods, GRAMPA, uses a distinct gene-tree reconciliation approach based on duplication and loss, but is restricted to inferring a single polyploidization event, limiting its applicability to datasets with more complex histories. We therefore developed GRAMPA^Iter^, an automated extension that recursively applies GRAMPA and combines the inferred events into a single MUL-tree, enabling its application to histories involving multiple polyploidization events. To characterize differences among the inferred networks, we further introduce two new measures that complement overall topological distance by separately quantifying disagreement in the descendant taxa and parental lineages associated with reticulation events.

## Materials and Methods

### Methods Evaluated

We evaluated five computational methods designed for polyploid phylogenetic inference: PADRE (Lott et al. 2009a), AlloppNET (Jones et al. 2013), GRAMPA (Thomas et al. 2017) together with our GRAMPA^Iter^ extension, MPAllopp (Yan et al. 2022), and Polyphest (Yan et al. 2024). The five methods differ in their inputs, models, and assumptions (Table 1; detailed descriptions in Supplementary Methods).

**Table 1.** Characteristics of the polyploid phylogeny reconstruction methods.

| Name (year) | Input / required prior information | Output | Description | Limitations |
|---|---|---|---|---|
| PADRE (2009) | Gene trees | MUL-tree, network | Builds a tree by first placing clades that can be placed in a single position, then remaining ones. Merges identical subtrees to get the network. | Does not model ILS. |
| AlloppNET (2013) | Sequence alignments and ploidy table | Posterior sample of MUL-trees | Bayesian method that jointly estimates the network, gene trees and assignment of tetraploid sequences to subgenomes under a model of ILS. | Restricted to diploid–tetraploid systems; requires ploidy information. |
| GRAMPA (2017) | Gene trees and singly labeled species tree | MUL-tree or singly labeled tree | Adapts reconciliation to MUL-trees to minimize duplication and loss. | Detects only a single event; does not model ILS. |
| MPAllopp (2022) | Gene trees | Network | Counts deep coalescences and searches heuristically for the most parsimonious network. | Heuristic search not guaranteed to be optimal. |
| Polyphest (2024) | Gene trees and ploidy set | MUL-tree, network | Extends PADRE to account for ILS by treating all clades equally and merging near-identical subtrees. | Requires ploidy information; performance depends on heuristically chosen thresholds. |

PADRE (Huber et al. 2006; Lott et al. 2009a, 2009b) was the first method designed specifically for reconstructing the evolutionary histories of polyploids. It first summarizes the gene trees into a consensus MUL-tree. Each gene tree is broken into its clades, and the clades are ranked by their frequency across gene trees. Clades that can be placed in only one position, because they contain a species represented by a single copy, are added to the MUL-tree first, followed by the remaining clades. Within each group, clades are added in order of frequency, and a clade is skipped if it cannot be placed in the MUL-tree together with those already added. PADRE then converts the MUL-tree into a network by merging identical subtrees in the MUL-tree, with each merge introducing a reticulation. Merging proceeds from the largest identical subtrees to the smallest.

AlloppNET (Jones et al. 2013; Jones 2017) is a Bayesian method that uses a probabilistic model of how ILS generates gene trees, evaluating candidate histories by how probable they make the observed data. Unlike the other methods considered here, it operates directly on sequence alignments rather than inferred gene trees, and requires an input table specifying whether each species is a diploid or tetraploid. It jointly estimates the network, the gene trees, and the assignment of each tetraploid’s sequences to its two parental subgenomes, sampling them from their posterior distribution. AlloppNET accommodates any number of species and hybridization events but is restricted to diploids and allotetraploids.

GRAMPA (Thomas et al. 2017) is based on gene-tree reconciliation, in which each gene tree is mapped onto a species tree to infer the minimum number of gene duplications and losses that explain it (Goodman et al. 1979; Page 1994), and extends it to MUL-trees. GRAMPA takes as input a collection of gene trees and a singly labeled species tree, and tests all possible pairs of nodes in the species tree, termed H1 and H2. H1 represents the polyploid clade, and H2 the second parental lineage. For each H1–H2 pair, the subtree rooted at H1 is copied and grafted onto the branch leading to H2, producing a candidate MUL-tree. All gene trees are reconciled against each candidate MUL-tree, and GRAMPA selects the MUL-tree with the lowest reconciliation score, defined as the total number of inferred gene duplications and losses across all gene trees. If H1 and H2 are placed on the same node, the event is interpreted as autopolyploidy, and otherwise as allopolyploidy. GRAMPA can detect at most one polyploidization event, a limitation we address with GRAMPA^Iter^ in the next section.

MPAllopp (Yan et al. 2022) also reconciles gene trees with MUL-trees, but its parsimony score counts deep coalescences, cases in which gene lineages fail to merge on the branch where they are expected to, as occurs under ILS. It extends the minimizing-deep-coalescences criterion (Maddison 1997) to polyploid networks. Its key modification for polyploids is that gene copies from different subgenomes are not expected to merge within the polyploid lineage, and so are not counted as deep coalescences. MPAllopp searches for the network with the fewest deep coalescences heuristically, and is therefore not guaranteed to find the optimal network.

Finally, Polyphest (Yan et al. 2024) builds on PADRE’s approach and modifies it to account for ILS. During consensus MUL-tree construction, Polyphest does not prioritize clades that can be placed in a single position. Instead, it selects the combination of clades with the highest total weight that can all be placed together in a single MUL-tree, where each clade is weighted by how many times it appears within gene trees, how often it appears across gene trees, and its size. During network construction, Polyphest relaxes PADRE’s requirement that merged subtrees be identical, allowing near-identical subtrees to be merged according to a user-defined threshold. Polyphest additionally requires prior information on the ploidy level of each taxon, thus constraining the expected number of subgenome copies in the output MUL-tree.

### GRAMPA^Iter^

To overcome GRAMPA’s limitation of inferring a single polyploidization event, Morales-Briones et al. (2021) inferred multiple events by manually pruning each inferred polyploid clade from the species and gene trees and re-applying GRAMPA to the remaining taxa. We developed GRAMPA^Iter^, an automated procedure that uses a recursive divide- and-conquer approach in which each GRAMPA-inferred polyploidization event partitions the current problem into two disjoint sub-problems. Events inferred across the resulting sub-problems are subsequently reassembled into a single MUL-tree.

At each recursion step, GRAMPA^Iter^ applies GRAMPA to identify the polyploidization event that minimizes the reconciliation score. The event is accepted only if the resulting MUL-tree has a lower reconciliation score than the input tree for that sub-problem. Upon acceptance, GRAMPA^Iter^ partitions the inferred MUL-tree into two new sub-problems: an inner subtree corresponding to the inferred polyploid clade, and an outer subtree corresponding to the MUL-tree with all copies of the polyploid clade removed. Importantly, each subtree is singly labeled. The gene trees are partitioned accordingly using the reconciliation mapping to the accepted MUL-tree. For each gene tree, each of the largest subclades that map exclusively to a single subgenome of the inferred polyploid lineage is assigned to the inner sub-problem as a separate gene tree, with the remainder assigned to the outer sub-problem. A sub-problem is discarded when its species tree is empty or when all of its gene trees have fewer than two leaves. The two sub-problems are then analyzed recursively, allowing the detection of multiple independent events including nested ones. The recursion terminates when no sub-problem yields an improved reconciliation score. This recursive partitioning produces a binary recursion tree of accepted events that records the hierarchy of the resulting sub-problems.

Reassembling the accepted events into a single global MUL-tree is non-trivial because each event’s recorded H2 location is defined relative to the local species tree present at that step of the recursion. During reassembly, the corresponding lineage may have been duplicated by another event, inferred elsewhere in the recursion, making its original H2 location ambiguous in the global MUL-tree. To resolve this ambiguity, GRAMPA^Iter^ traverses the recursion tree in post-order and recreates each accepted event by grafting the inner sub-tree onto the outer sub-tree at its original position and the recorded H2 location of each of its copies. In the partially reconstructed MUL-tree, when H2 maps to multiple positions corresponding to identical sister clades (as in autopolyploidy) or identical paraphyletic groups (as in allopolyploidy along one branch), GRAMPA^Iter^ identifies the lowest nodes whose children are exclusively matching positions (i.e., H2 nodes, or original) and places a single graft at that position, rather than grafting separately onto each copy, thereby preserving the hierarchy of events (Fig. 2). Otherwise, a graft is inserted at each H2-mapping branch.

**Figure 2.**
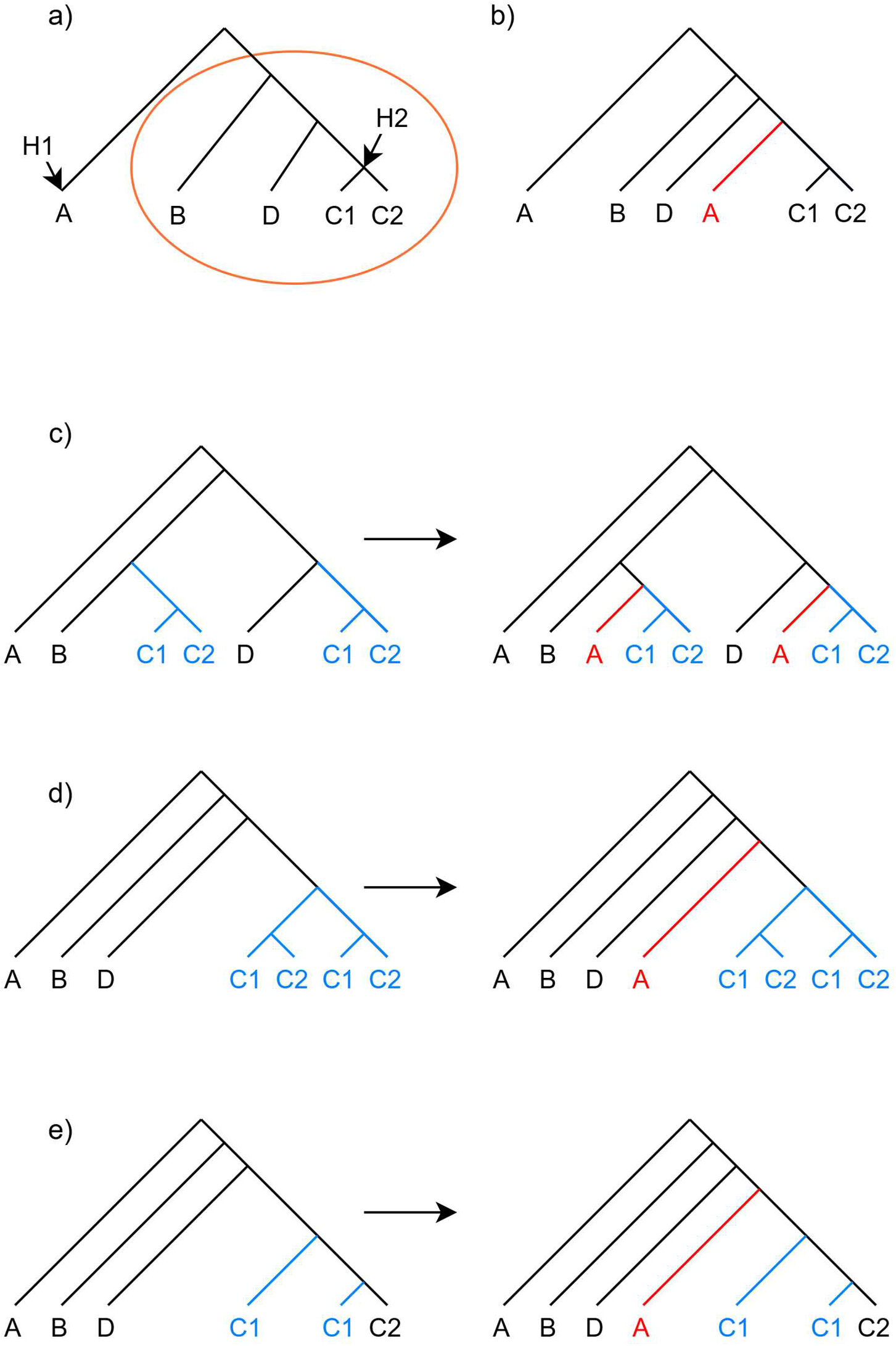
Reassembly of an accepted polyploidization event in GRAMPA^Iter^. (a) Local species tree at the recursion step where a polyploidization event is inferred, with H1 = A and H2 = the node leading to clade (C1, C2). The event duplicates the subtree rooted at H1 and grafts one copy onto H2. The orange circle marks the outer subtree, which may have been modified by other events by the time reassembly occurs. (b) Standard reassembly when no subsequent event has modified H2 such that a second copy of A is grafted at the recorded H2 location. (c–e) Reassembly when an event reconstructed earlier during reassembly has created multiple mappings of H2. In each panel, the left side shows the partially reconstructed MUL-tree before incorporation of the focal event and the right side shows the result after its incorporation. (c) Separate grafts: when the H2 mappings occur at separate positions, A is grafted independently onto each. (d) Single graft: When H2 maps to two identical sister clades, as in the autopolyploidy case shown here, a single copy of A is grafted above their common ancestor. (e) Single graft: a previously reassembled event has duplicated C1 along the branch leading to clade (C1, C2), so H2 maps to the two copies of C1, which do not form a clade. In this case, a single copy of A is grafted at the lowest position ancestral to all H2 mappings. In all panels, black branches show the original tree, blue branches indicate lineages duplicated by previously reassembled events, and red branches show the copies of A grafted during reassembly.

In the new implementation, we also increased the “group cap” parameter, which GRAMPA uses to filter excessively complex gene trees by limiting the number of ambiguous gene-copy mappings considered during reconciliation. This was enabled by a more efficient implementation of the code, allowing us to raise the cap from 8 to 15 and thereby retain gene trees with more complex polyploid signal that would otherwise have been excluded. For consistency, the results of GRAMPA reported here correspond to the first iteration of GRAMPA^Iter^, rather than separate runs of the original GRAMPA package, and therefore also use a group cap of 15.

### Running the Methods

All methods were applied to both the empirical and simulated datasets with a runtime limit of five days per analysis. GRAMPA, GRAMPA^Iter^ and AlloppNET, through BEAST, support multithreading, and these were run with eight CPU cores. The remaining methods are single-threaded. PADRE was run using its command-line version 2.0. GRAMPA v1.4.1 and GRAMPA^Iter^ require a singly labeled species tree as input, which was inferred from the gene trees using ASTRAL IV as implemented in ASTER v1.23 (Zhang et al. 2025). When gene trees were unrooted, they were rooted, and polytomies resolved, using NOTUNG v2.9.1.5 (Stolzer et al. 2012). Polyphest was run with a default near-identical subtree threshold of 0.2, and with clade-support percentile thresholds of 50%, 70% and 90%, retaining clades whose support exceeded the corresponding percentile of the support distribution within each dataset. For each dataset, we report the result from the lowest threshold at which the analysis completed, following the authors’ recommendation (Z. Yan and L. Nakhleh, personal communication). MPAllopp was run through MP-SUGAR, its Python implementation in the PhyNetPy package v0.0.8, as recommended by its authors (Z. Yan and L. Nakhleh, personal communication). AlloppNET was run in BEAST v1.8.2 (Drummond and Rambaut 2007) with a target chain length of 10^8^ iterations. Because no chain reached this length within the runtime limit, the sampled MUL-trees were summarized as a maximum clade credibility tree using TreeAnnotator v1.8.2 (Drummond and Rambaut 2007), after discarding the first 10% of samples as burn-in. Polyphest and AlloppNET require the ploidy of each species as input. Unless reported in the original study, ploidy was inferred from gene-copy numbers as described in the Empirical Analysis section. Inferred networks were visualized with Dendroscope v3.8.10 (Huson and Scornavacca 2012), which converts MUL-trees into networks using the same algorithm as PADRE’s second step (Huber et al. 2006).

### Network Comparison Measures

We used three measures to evaluate the similarity between two networks: two measures that compare the inferred reticulations, and the μ-distance, which measures overall topological disagreement between the networks. The evaluated methods differ in the representation of their output, returning either a MUL-tree or a phylogenetic network. Comparisons in this study are made between networks, so every output was brought to that representation before being scored. Unfolding a network into its corresponding MUL-tree is deterministic, whereas folding a MUL-tree into a network can depend on the algorithm used. We therefore folded every inferred MUL-tree into a network using our implementation of the HOLM algorithm (Huber et al. 2006). Outputs already returned as a network were first unfolded to their MUL-tree and then folded again by the same procedure. This common folding procedure ensured that differences among the resulting networks were not introduced by the conversion algorithm.

Because reticulations in different networks are neither labeled nor ordered, correspondence between events was first inferred as follows. For every pair of reticulations, one from network A and one from network B, we computed the Jaccard similarity between their sets of descendant leaves and obtained the one-to-one matching that maximizes the total Jaccard similarity by applying the Hungarian algorithm (Kuhn 1955), as implemented in SciPy’s linear_sum_assignment (Virtanen et al. 2020). This matching was used to compute two measures. The reticulation descendants measure compares the leaf sets of each matched pair of reticulation nodes, regardless of whether the event is an auto- or an allopolyploidization.

The reticulation sisters measure compares the parental context of each matched pair. For each parent of a reticulation node, the sister clade is defined as the set of descendant species excluding those reached through the reticulation itself. Autopolyploidization events share the same sister clade, while allopolyploidization events have one sister clade per parental lineage. Within each matched pair, sister clades were paired using a second Hungarian assignment. Jaccard distances were summed across matched sister clades, with unpaired clades contributing 1.0, and divided by the larger number of sister clades.

For both measures, a Jaccard distance is computed for each matched pair, unmatched reticulation nodes in the larger set each contribute the maximum distance of 1.0, and the total is normalized by *max*(*m*, *n*). Both therefore range from 0, when all reticulations match with identical sets, to 1, when no overlap exists. We additionally recorded the signed difference in the number of reticulation events between the two networks, where a reticulation node with more than two parents contributes one event per additional parent.

To quantify global topological disagreement between two networks, we computed the μ-distance. In the classical μ-distance, each node is described by a vector counting the paths that lead from it down to each species, and the network is described by the collection of these vectors (Cardona et al. 2009). Because folding can result in a reticulation node with more than two parents, these vectors no longer identify a network uniquely. Therefore, we appended the in-degree of each node to its vector, following Reichling et al. (2026). The μ-distance is defined only for networks on the same taxon set. The distance between two networks is then the number of these vectors present in one network but absent in the other. Leaves are excluded because their vectors are identical across networks with the same taxon set. The resulting count is normalized by the total number of internal nodes across both networks. Values range from 0 for isomorphic networks to 1 when the two share no internal-node vector. The μ-distance is a valid metric only on a restricted class of networks (Reichling et al. 2026), which includes 14 of the 21 model networks used here. We computed the vectors with the phylonetwork Python package version 2.2.2 (Cardona et al. 2024) and added the parent counts using a Python script.

Differences between methods were tested with two-sided Wilcoxon signed-rank tests (Wilcoxon 1945). For the simulations, replicate values were first averaged per network, and tests were paired by network within each configuration. To test whether one method was more sensitive than another to a simulation parameter, we fitted a least-squares line to each measure against the parameter level for each network and method, and compared the slopes of the two methods with a paired Wilcoxon signed-rank test. Simulation p-values were corrected for multiple testing with the Holm– Bonferroni procedure (Holm 1979) separately for each comparison measure within each analysis. For example, the comparison of Polyphest and GRAMPA^Iter^ in the baseline simulations was corrected across the 12 configurations. For the empirical data, tests were paired by dataset. Relationships among the comparison measures across empirical datasets were assessed using Spearman rank correlations. All tests used SciPy v1.12.0 (Virtanen et al. 2020).

### Empirical Analysis

We conducted a literature survey to identify published studies reporting phylogenetic networks that included at least one polyploid species and for which gene trees or gene sequence alignments were available. Studies providing only raw sequencing reads were excluded. The survey identified 21 datasets spanning diverse plant lineages, plus one alga, and ploidy levels, from diploid–tetraploid systems to complexes containing taxa with ploidy levels as high as 18x. The number of genes ranged from a single gene in nine datasets to over 4,000 loci. When gene trees were available, they were used directly; otherwise, gene trees were inferred from the available alignments using IQ-TREE v3.0.1 (Wong et al. 2026). The 21 datasets and their main characteristics are summarized in Supplementary Table S1.

Because the original studies applied diverse preprocessing pipelines and the available data were obtained at different stages of these pipelines, their exact preprocessing procedures could not be reproduced consistently. We therefore applied a standardized gene-copy selection procedure across datasets. In many gene trees, taxa were represented by more gene copies than the number of subgenomes represented in the published network. For each taxon, we retained up to *k* copies, where *k* is the expected number of subgenomes based on the known or inferred ploidy level (*k* = 1 for diploids, *k* = 2 for tetraploids, and so on). When a gene tree contained *k* or fewer copies, all were kept. When more than *k* copies were present, copies were selected greedily using a Python script. When a single copy was retained, the copy most distant from the root was kept. Otherwise, selection started from the two most distant copies and then repeatedly added the copy whose distance to the nearest already selected copy was largest. Distances were computed from branch lengths when available and otherwise as the number of branches separating copies in the gene tree. Copies were selected independently for each taxon. This strategy favors the retention of divergent copies that may represent distinct subgenomes, regardless of whether copies form monophyletic groups or are dispersed across the gene tree.

Where ploidy was unknown, we inferred it from the number of gene copies observed for each taxon. For each taxon, we counted its copies in every gene tree in which it was present, yielding a distribution of copy numbers across gene trees. Because gene duplication and loss cause copy number to vary among genes, the most frequent copy number does not necessarily reflect the number of subgenomes. We therefore smoothed this distribution before selecting a value. Each candidate copy number received a score equal to the number of gene trees with that copy number, plus the number of gene trees with neighboring copy numbers, weighted according to a triangular kernel of width two (i.e., placing weight of 2/3 for copy numbers differing by one and weight of 1/3 for those differing by two). The copy number with the highest score was taken as the number of subgenomes. For example, a taxon observed with one copy in three gene trees, two copies in two, and three copies in two would be assigned one copy based on the most frequent value alone, but two copies after smoothing, because the combined evidence from two and three copies outweighs that for a single copy. When a taxon showed only one or two distinct copy numbers, the more frequent one was used directly.

Because AlloppNET is restricted to diploid and tetraploid taxa, it was applied only to four datasets in which the inferred maximum copy number was two. Pairwise comparisons were restricted to the 12 empirical datasets containing multiple gene trees. When a method failed to produce a result for a dataset, pairwise comparisons involving that method were omitted for that dataset.

### Simulation Study

To enable assessment of method accuracy, we conducted simulations in which the ground-truth evolutionary history was known. To obtain networks reflecting the size, ploidy levels and reticulation structure of real polyploid systems, we assembled 21 MUL-trees derived from the empirical datasets (see Empirical Analysis; Supplementary Table S1). These model networks comprised 5 to 82 taxa (median 18) and 1 to 49 polyploidization events (median 8; Supplementary Table S2). For 16 datasets, the model MUL-tree was either obtained directly from the original study or, for single-gene datasets where only sequences were available, inferred from those sequences using IQ-TREE v3.0.1 (Wong et al. 2026). For the remaining five multi-gene datasets, for which no published MUL-tree was available, we used a MUL-tree inferred by one of the methods evaluated here, as specified in Supplementary Table S2. Because these trees served only as model topologies and the gene trees were subsequently simulated independently, this choice did not directly favor the method used to infer the model tree. All 21 MUL-trees were rescaled to a root-to-tip height of 10 million generations.

Our base simulation design comprised 12 configurations organized as a 3 × 4 grid (Table 2), crossing three ILS levels (low, medium, high) with four duplication and loss levels (none, low, medium, high). For each configuration, we simulated 1,000 gene trees per network with five replicates, yielding 105 datasets per configuration and 1,260 datasets overall. Gene trees were simulated using SimPhy v1.0.2 (Mallo et al. 2016). SimPhy requires singly labeled species trees. Each model MUL-tree was therefore converted by appending numeric suffixes to duplicated labels, so that a species *X* appearing *K* times became *X*_1_, …, *X*_K_. SimPhy simulates duplication and loss of each gene family along this relabeled species tree, producing a locus tree, and then simulates gene trees within each locus tree under the multispecies coalescent. In our simulations, duplication and loss rates and the effective population size were constant across lineages and loci, and horizontal gene transfer, gene conversion and selection were not included. The suffixes were removed after gene tree generation, restoring the original species names. ILS levels were set by varying the effective population size (Ne; Table 2). To quantify the resulting discordance, we computed the normalized Robinson– Foulds distance (Robinson and Foulds 1981) between each simulated gene tree and the species tree from which it was simulated. Because gene duplication and loss change the set of leaves in a gene tree, nRF was computed only in the three configurations without duplication and loss, in which every gene tree contains exactly one copy of each species-tree leaf. Averaged across gene trees, replicates and networks, this yielded mean nRF values of approximately 1% (low, Ne = 200,000), 27% (medium, Ne = 1,000,000) and 49% (high, Ne = 2,000,000). These levels span from near-complete agreement to substantial discordance and follow ranges used in previous studies (Sayyari and Mirarab 2016; Willson et al. 2022; Tabatabaee et al. 2025). Gene duplication and loss rates spanned empirical estimates across diverse plant lineages (Lynch and Conery 2003; Guo 2013; Wang et al. 2018), and were set to none, 10^−9^, 10^−8^ and 10^−7^ per gene per generation. Together, these processes are referred to here as gene turnover.

**Table 2.**
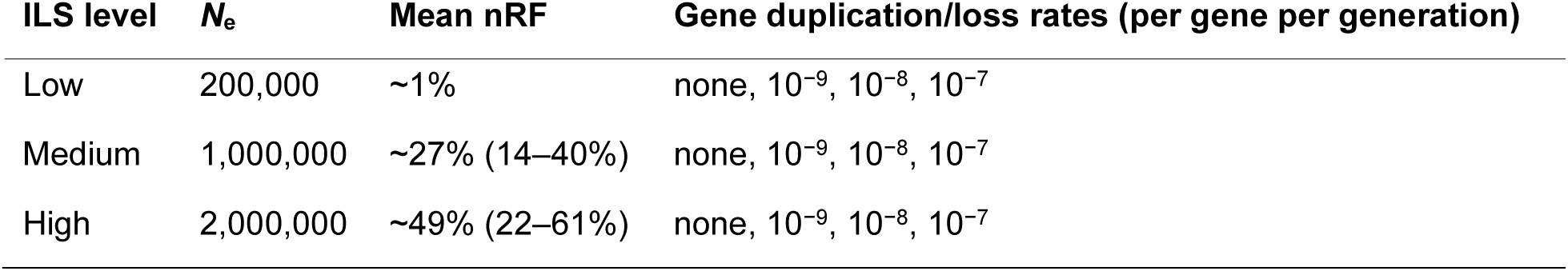
The 12 baseline simulation configurations, crossing three ILS levels with four gene duplication and loss rates. . nRF: mean normalized Robinson–Foulds distance between the simulated gene trees and the species tree, computed in the configurations without gene duplication and loss. Values in parentheses give the range across the 21 networks.

| **ILS level** | ***N*<sub>e</sub>** | **Mean nRF** | **Gene duplication/loss rates (per gene per generation)** |
|---|---|---|---|
| Low | 200,000 | ~1% | none, 10<sup>−9</sup>, 10<sup>−8</sup>, 10<sup>−7</sup> |
| Medium | 1,000,000 | ~27% (14–40%) | none, 10<sup>−9</sup>, 10<sup>−8</sup>, 10<sup>−7</sup> |
| High | 2,000,000 | ~49% (22–61%) | none, 10<sup>−9</sup>, 10<sup>−8</sup>, 10<sup>−7</sup> |

For AlloppNET, which requires sequence data, DNA sequences were simulated along the gene trees using AliSim (Ly-Trong et al. 2022) as implemented in IQ-TREE v3.0.1 (Wong et al. 2026), under the GTR+Γ substitution model, with alignment lengths, substitution rates, GTR rates, base frequencies and the gamma shape parameter sampled jointly from distributions inferred from 2,709 gene alignments across three data-rich empirical datasets.

For all other methods, we supplied the SimPhy simulated gene trees directly rather than re-inferring them from the simulated sequences. This isolated the performance of network inference from the accuracy of gene tree estimation. Although the true ploidy levels and copy numbers were known from the ground truth networks, we did not provide this information to any method. Instead, methods requiring ploidy information received inferred inputs with the kernel smoothing approach described in the Empirical Analysis section, so that the simulation reflected realistic conditions in which true ploidy levels are unknown.

Each method was run separately on every simulated dataset, defined by one network, one replicate and one configuration. All methods were applied to all 105 datasets of each configuration, except AlloppNET, which cannot analyze taxa above tetraploid level (see Methods Evaluated) and was therefore applied only to the 40 datasets per configuration derived from the eight networks containing only diploid and tetraploid taxa. Accuracy was quantified against the true networks using the three measures (see Network Comparison Measures): the reticulation descendants measure, the reticulation sisters measure, and the µ-distance. All measures were computed independently for each replicate and aggregated as the mean and standard error.

#### Simulations accounting for post-WGD diploidization

—The base simulations detailed above model gene duplication and loss as a continuous birth-death process but do not capture the extensive loss of duplicated gene copies that typically follows WGD during diploidization (Freeling 2009; Qiao et al. 2019). We therefore conducted an additional simulation experiment in which this post-WGD gene loss, termed fractionation, was applied to the simulated gene trees after they were generated. Applying it required knowing which gene copies were produced by each WGD event. The model MUL-trees specify the subgenome copies of each polyploid taxon but not the events that produced them. We therefore identified the events and the set of copies each produced by folding each MUL-tree with the HOLM algorithm. Because the copy suffixes added for SimPhy (*X*_1, …, *X*_*K*) were still present in the simulated gene trees at this stage, each gene copy could be assigned directly to the event that produced it.

Following Rabier et al. (2014), each event was assigned a fixed per-event retention probability *q*, representing the expected fraction of duplicated genes that retain both copies. For each gene tree and each taxon, WGD events were processed from oldest to most recent, keeping one copy at random and removing each of the other copies with probability 1 − *q*. Because fractionation acts on the existing gene trees, all retention settings share the same underlying simulation. Fractionation was applied to the base configuration of medium-ILS (Ne = 1,000,000) and medium-duplication/loss rates at three retention probabilities of *q* = 0.75, 0.50, and 0.25, spanning most of the empirical range reported across WGDs (Tiley et al. 2016). Fractionation was not applied to the network of Morales-Briones et al. (2021), on which the methods did not terminate in most replicates of the base configuration. The corresponding unfractionated base configuration (*q* = 1) served as the reference, yielding a four-level retention series from no fractionation to extensive gene loss.

## Results

### A Framework for Comparing Inferred Networks

Our primary interest was whether a method recovers the polyploidization events themselves, namely which lineages are polyploid and the parental lineages associated with each polyploidization event. We therefore introduced two reticulation-specific measures that address these components directly, while the μ-distance summarizes structural disagreement over the whole network. This comparison framework distinguishes among types of disagreement that are conflated by a single global score. As a minimal illustrative example (Fig. 3), the reference network places polyploid species *p* as a hybrid of *q* and *r*. We compared three inferred networks against it, each introducing a single type of error. The two reticulation measures resolve these errors into distinct signatures (Table 3). In Figure 3b, the wrong taxon is inferred as polyploid (*u* instead of *p*). This error is detected entirely by the reticulation descendants measure, which reaches 1.0 because the lineage descending from the reticulation is *u* rather than *p*, while the reticulation sisters measure is 0 because the parental clades are identical. In Figure 3c, one parental lineage is incorrectly inferred (*s* instead of *r*). Here, the reticulation sisters measure is 0.5, while the reticulation descendants measure is 0. In Figure 3d, a spurious second reticulation involving *u* is inferred. In this case, both measures are 0.5, reflecting that one inferred reticulation matches the reference perfectly while the second has no counterpart. Each error type therefore produces a characteristic signature.

**Figure 3.**
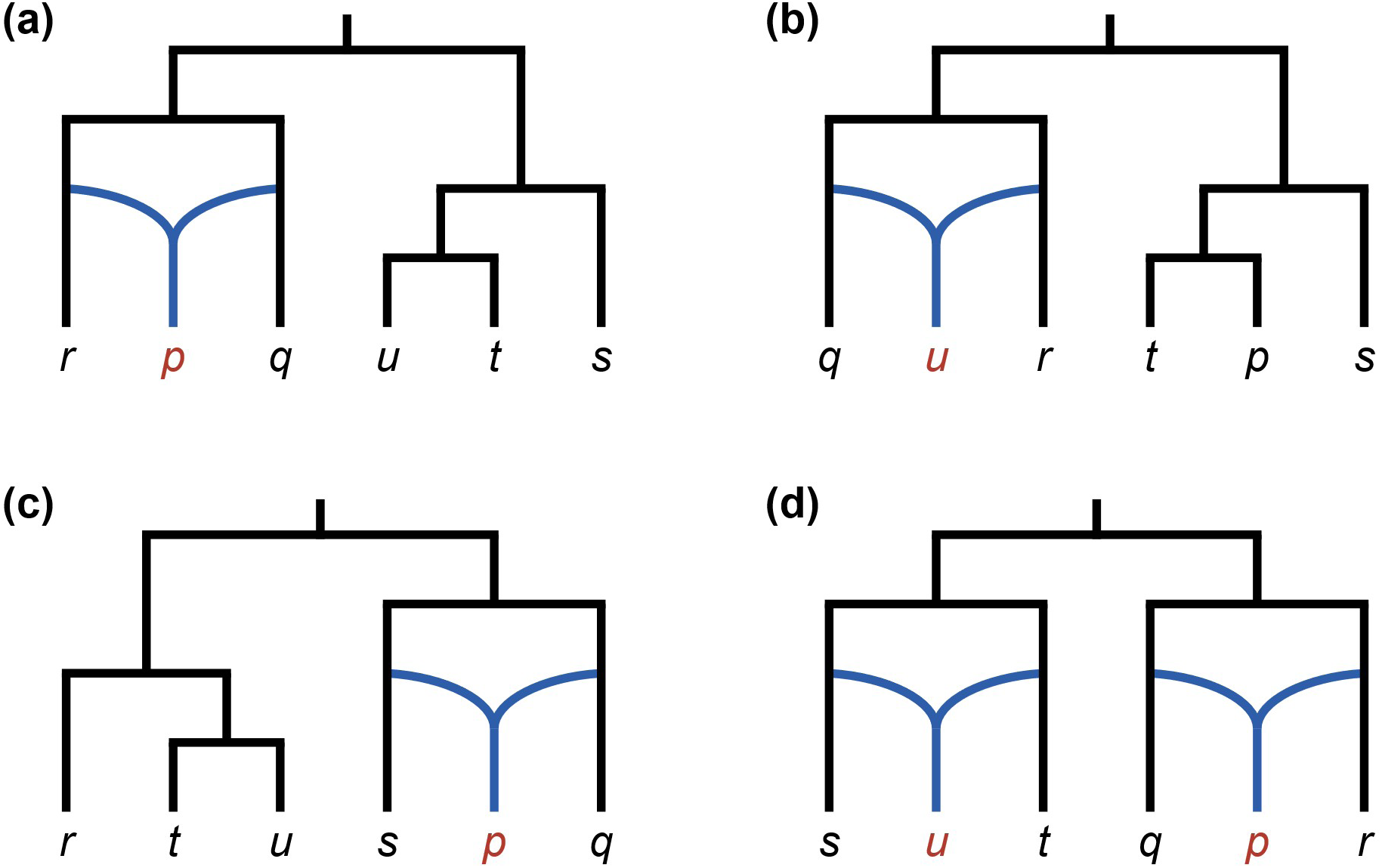
Demonstration of the comparison framework on an illustrative example. The true network in panel (a) represents polyploid species p as a hybrid of q and r. Each of the three inferred networks introduces a different error. (b) Wrong polyploid species: the parental lineages are correct, but u rather than p is inferred as polyploid. (c) Wrong parental lineage: p is correctly inferred as polyploid but as a hybrid of q and s instead of q and r. (d) Extra reticulation: p and its parental lineages are correctly inferred, but an additional reticulation erroneously identifies u as polyploid. The effects of these three error types on the comparison measures are summarized in Table 3.

**Table 3.** Comparison measures for the three error types illustrated in Figure 3. Values show the disagreement between the true network (Fig. 3a) and networks containing a wrong polyploid species *(Fig. 3b), a wrong* parental lineage *(Fig. 3c), or an extra reticulation (Fig. 3d), with 0 indicating complete agreement*.

| **Comparison measure** | **Wrong polyploid species** | **Wrong parental lineage** | **Extra reticulation** |
|---|---|---|---|
| Reticulation descendants measure | 1.0 | 0 | 0.5 |
| Reticulation sisters measure | 0 | 0.5 | 0.5 |
| Difference in reticulation count (signed) | 0 | 0 | +1 |
| μ-distance | 1.0 | 0.43 | 0.38 |

The μ-distance is nonzero in all three cases, confirming that each inferred network differs from the reference, but provides no indication regarding the source of disagreement. It assigns similar values to the wrong-parent and the extra-reticulation errors (0.43 and 0.38, Figs. 3c, 3d), despite their fundamentally different nature. On the other hand, the μ-distance reaches its maximum of 1.0 for the incorrectly inferred polyploid species (Fig. 3b), because the two networks share no internal-node vectors. This case thus illustrates saturation of the μ-distance, which cannot distinguish this error from networks showing even greater structural disagreement. Moreover, because the μ-distance measures disagreement across the entire network topology, differences in the placement of diploid species contribute to the score similarly to differences in the polyploidization events themselves. We therefore report the μ-distance as a summary of overall network disagreement, while relying on the reticulation-specific measures to characterize disagreement in the inferred polyploidization events.

### Wide Disagreement between Inference Methods on Empirical Data

We applied the five polyploid phylogenetic inference methods to 12 empirical datasets obtained from the literature. The methods differed substantially in completion rate, in which species they inferred to be polyploid, in the number and placement of reticulations, and in overall network topology. GRAMPA and GRAMPA^Iter^ completed the analyses on all 12 datasets, Polyphest on 11, PADRE on two, and MPAllopp on one. AlloppNET was applied to only four datasets due to its restriction to diploid-tetraploid systems. Runtimes differed by orders of magnitude: GRAMPA^Iter^ completed in under 65 seconds on all datasets, whereas Polyphest ranged from under one second to over 35 hours. MPAllopp reached the five-day limit on all but one dataset. AlloppNET reached this limit on all four applicable datasets, and so its reported results represent chains that were terminated before convergence. Because GRAMPA is restricted to a single event and only Polyphest and GRAMPA^Iter^ produced multi-event inferences for most datasets, the broader analysis focused primarily on these two.

#### Two illustrative datasets

*—*The dataset of Popp et al. (2005) represents a relatively simple example, comprising six low-copy nuclear genes from 14 species of *Silene* section *Physolychnis*. The original study did not apply a network inference method. Instead, it reconstructed the history manually by comparing the gene trees and identifying distinct homeologs derived from different parental lineages in the polyploids. It proposed two consecutive allopolyploidization events: *S. uralensis* and *S. ajanensis* hybridized to produce the allotetraploid lineage leading to *S. involucrata*, which subsequently hybridized with an additional lineage to produce the hexaploids *S. sorensenis* and *S. ostenfeldii*. Because the original study presented a network of only eight of the 14 species, our measures could not be used to compare it with the inferred networks, and we therefore describe their agreement qualitatively. Polyphest received as input the ploidy levels reported in the original study, while the other methods did not. AlloppNET was not applicable and MPAllopp failed to complete within the runtime limit. The three methods that completed the analysis inferred substantially different networks (Supplementary Figs. S1, S2), with Polyphest inferring five reticulations, GRAMPA^Iter^ seven, and PADRE four. PADRE returned a network containing 13 of the 14 analyzed species, omitting *S. linnaeana*, the species represented in the fewest gene trees.

All three methods identified a hybridization event leading exclusively to *S. involucrata*, in agreement with the original study, but differed in its inferred parental lineages, the surrounding topology, and the additional events inferred. Polyphest and PADRE both recovered the three polyploid species proposed in the original study, each associated with a distinct reticulation node leading to a single descendant, so the reticulation descendants measure between the two networks was 0. For Polyphest, this inference was constrained by the ploidy information it received as input, whereas PADRE recovered it without such prior data. The larger numbers of reticulations reported above reflect reticulation nodes with more than two parents, each representing multiple events. The two networks disagreed on the origins of *S. involucrata* and *S. sorensenis*. Polyphest inferred *S. involucrata* as an allotetraploid, as in the original study, and *S. sorensenis* as a hexaploid formed by autopolyploidization followed by hybridization. PADRE inferred both species as autotetraploids, contrary to the allopolyploid origins proposed in the original study. These differences were reflected in a high reticulation sisters measure between the Polyphest and PADRE networks (0.62). GRAMPA^Iter^ inferred a different scenario. After identifying the *S. involucrata* event in its first iteration, it continued to infer additional reticulations, eventually recovering seven events involving ten polyploid species.

The dataset of Wu et al. (2016) provides a more complex example, comprising two low-copy nuclear gene trees from 18 taxa of *Ephedra*, for which flow cytometry identified nine tetraploids and nine diploids. These assignments were supplied as priors to Polyphest and AlloppNET. The original study inferred its network with PADRE, selecting gene copies manually based on their positions in the gene trees. Our PADRE network, based on our standardized gene-copy selection, identified the same polyploid taxa as the published network but differed in some of the inferred parental lineages. Polyphest, PADRE, and AlloppNET all inferred networks with the same set of nine polyploid taxa, consistent with the flow cytometry data. For Polyphest and AlloppNET,

this agreement was imposed by the ploidy prior, whereas PADRE recovered the same set independently. GRAMPA^Iter^ instead identified 16 of the 18 taxa as polyploids, all resulting from one autopolyploidization event inferred at the base of the phylogeny in the first inference iteration. Five additional reticulations subsequently led to six taxa inferred as octoploid (Fig. 4b).

**Figure 4.**
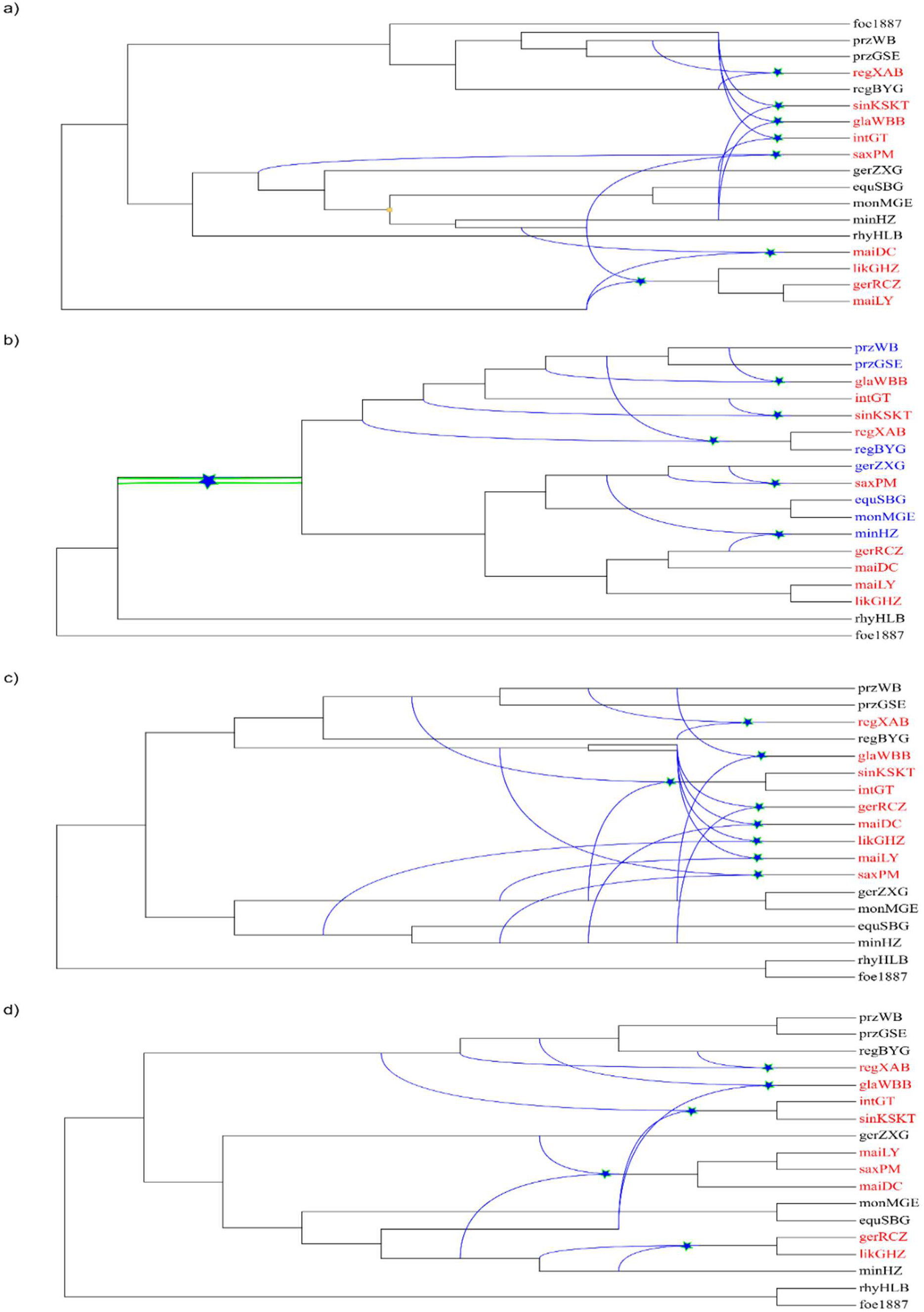
Inferred networks for the Ephedra dataset of Wu et al. (2016). Red taxa indicate the nine tetraploid species identified in the original study and provided as prior information to Polyphest and AlloppNET. Blue taxa indicate additional species inferred as polyploid by GRAMPA^Iter^. Black taxa indicate species inferred as diploid in the corresponding network. Stars denote reticulation nodes representing WGD events (autopolyploidization or allopolyploidization). (a) Polyphest (7 events, 9 polyploids). (b) GRAMPA^Iter^ (6 events, 16 polyploids). The green double edge marks the autopolyploidy event inferred by GRAMPA^Iter^ in its first iteration. (c) PADRE (8 events, 9 polyploids), the method used to infer the network in the original study. (d) AlloppNET (5 events, 9 polyploids).

Crucially, agreement on which species are polyploid did not translate into agreement on where the reticulations were placed. Agreement on reticulation descendants was greatest between Polyphest and PADRE (score = 0.40), followed by Polyphest and GRAMPA^Iter^ (0.47). Despite receiving identical ploidy priors, AlloppNET and Polyphest showed greater disagreement, with a reticulation descendants score of 0.50. Disagreement on the parental lineages was greater still, and for every pair of inferred networks the reticulation sisters measure exceeded the reticulation descendants measure, ranging from 0.63 between Polyphest and PADRE to 0.85 between PADRE and GRAMPA^Iter^ and between Polyphest and AlloppNET. The µ-distance also placed Polyphest and PADRE closest, at 0.53, but ordered the remaining pairs differently: it ranked PADRE and GRAMPA^Iter^ third closest, at 0.73, although these two agreed least of all on the inferred reticulation descendants (0.62). It gave Polyphest and AlloppNET the same value as Polyphest and GRAMPA^Iter^ (µ-distance = 0.79), so sharing the same ploidy prior did not result in greater similarity between the Polyphest and AlloppNET networks. Visual inspection confirms substantial topological differences: Polyphest and PADRE assigned individual polyploid species to separate, independent reticulations, whereas GRAMPA^Iter^ and AlloppNET inferred some reticulations at deeper nodes corresponding to larger polyploid clades. No reticulation event was consistently recovered across all methods.

#### Comparison of Polyphest and GRAMPA^Iter^ across all empirical datasets

*—*Across all 11 datasets for which both Polyphest and GRAMPA^Iter^ produced results, disagreement on the inferred reticulations was high. The reticulation descendants measure ranged from 0.33 to 0.96, with a mean of 0.73, while disagreement over parental lineages was higher still, with the reticulation sisters measure ranging from 0.72 to 1.00 (mean = 0.87) and reaching complete disagreement for the dataset of Ding et al. (2023). The two measures were strongly correlated across datasets (Spearman ρ = 0.79, *p* = 0.004), so datasets on which the methods disagreed on the descendant taxa also tended to show disagreement on the parental lineages.

GRAMPA^Iter^ inferred significantly more polyploid species than Polyphest on all 11 datasets, with a mean of 18.6 compared with 9.9 (Wilcoxon signed-rank test, *p* = 0.001). In contrast, GRAMPA^Iter^ inferred fewer reticulations, with means of 6.9 compared with 12.7 for Polyphest, although this difference was statistically non-significant (*p* = 0.08). The larger polyploid sets inferred by GRAMPA^Iter^ reflected where its reticulations were placed. Reticulations inferred by GRAMPA^Iter^ had an average of 4.8 descendant species, compared with 1.0 for Polyphest (*p* = 0.001), which almost always assigned each reticulation to a single species. GRAMPA^Iter^ thus placed reticulations at deeper nodes, where a single event rendered an entire clade polyploid. In the dataset of Morales-Briones et al. (2021), for example, GRAMPA^Iter^ identified 73 polyploid species compared with 49 for Polyphest, despite inferring fewer reticulations (18 versus 49).

The µ-distance between the two methods ranged from 0.53 to 0.90, with a mean of 0.72, so even on the most similar dataset more than half of the internal-node vectors were unmatched between the two networks. Across the 11 datasets, both correlations between the µ-distance and the reticulation-specific measures were negative but non-significant (reticulation descendants: Spearman ρ = −0.50, *p* = 0.12; reticulation sisters: Spearman ρ = −0.26, p = 0.43). The dataset on which the two methods were closest by µ-distance, that of Wisecaver et al. (2023) at 0.53, was among those on which they disagreed most on the inferred reticulations (reticulation descendants 0.92, reticulation sisters 0.95). Conversely, both reticulation measures were lowest on the dataset of Ren et al. (2024), at 0.33 and 0.72, which ranked only fourth by µ-distance.

### Accuracy on Simulated Data

*Completion rates.—*To evaluate method performance, we simulated five replicate datasets for each of 21 model networks under 12 configurations, combining three levels of ILS with no, low, medium or high gene turnover rates, resulting in 105 runs per method and configuration (see Simulation Study). Completion rates varied substantially across methods and configurations (Fig. 5). GRAMPA completed all 105 runs in every configuration, while GRAMPA^Iter^ completed 94–96%, with nearly all failures confined to the network of Morales-Briones et al. (2021), the most complex model network (comprising 82 species and 49 polyploidization events), on which the iterative procedure frequently did not terminate. For these failed inferences, the first-iteration GRAMPA results were retained for subsequent accuracy analyses. Polyphest completed 76–85% of runs across all configurations. The remaining methods completed far fewer runs within the time limit. PADRE completed only 10% and 4% of runs under low and medium ILS, respectively, in the absence of gene turnover, and failed entirely under high ILS and under every configuration involving gene turnover.

**Figure 5.**
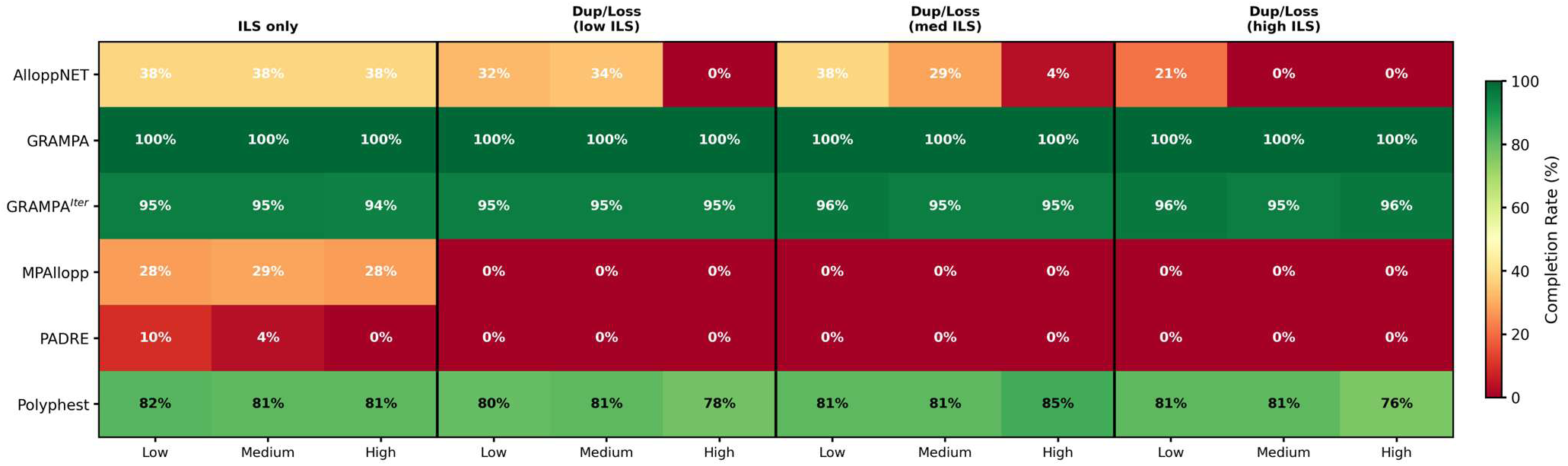
Completion rates of all methods across the 12 baseline ILS × gene duplication/loss simulation configurations without post-WGD fractionation. Each cell shows the percentage of completed replicates, with darker shading indicating higher completion.

MPAllopp completed 28–29% of runs across the three ILS-only configurations but failed entirely whenever gene turnover was included. AlloppNET, restricted to the eight tetraploid networks, completed 29–38% of runs under low to medium rates of gene turnover, dropping to 0–10% under high rates. Given these patterns, subsequent accuracy analyses focus primarily on Polyphest and GRAMPA^Iter^, the only multi-event methods that consistently produced results, with the remaining methods assessed separately on the subset of eight networks containing only diploid and tetraploid taxa.

#### Polyphest was more accurate than GRAMPA^Iter^ across all baseline configurations

*—*Polyphest outperformed GRAMPA^Iter^ across all 12 baseline configurations (Fig. 6). Under the simplest setting of low ILS and no gene duplication and loss, Polyphest achieved a mean reticulation descendants measure of 0.03 against 0.48 for GRAMPA^Iter^ and a mean reticulation sisters measure of 0.06 against 0.65. The μ-distance showed the same ordering, 0.03 against 0.52. Thus, under the simplest conditions, Polyphest showed near-perfect agreement with the true networks in both the descendant taxa and parental lineages, whereas GRAMPA^Iter^ showed substantial disagreement on both components. Polyphest had significantly lower μ-distance and reticulation sisters measures in all 12 configurations, and a significantly lower reticulation descendants measure in 10 of them (Holm-adjusted p < 0.05). Accuracy of both methods decreased with increasing ILS and gene turnover (Fig. 6), although the effect of gene turnover was generally more pronounced. Polyphest was particularly sensitive to increasing gene turnover, whereas GRAMPA^Iter^ showed a more moderate response. The difference between the methods narrowed most strongly at high rates of gene turnover: at low ILS, Polyphest’s reticulation descendants measure increased from 0.03 to 0.33 across gene-turnover gradients, whereas GRAMPA^Iter^’s increased only from 0.48 to 0.53. The two methods also differed in their estimates of the number of reticulations. Polyphest was close to the true number of reticulations under low ILS with no or moderate gene turnover, but increasingly overestimated their numbers as ILS or gene turnover intensified. GRAMPA^Iter^ showed the opposite tendency, underestimating the number of reticulations in every configuration.

**Figure 6.**
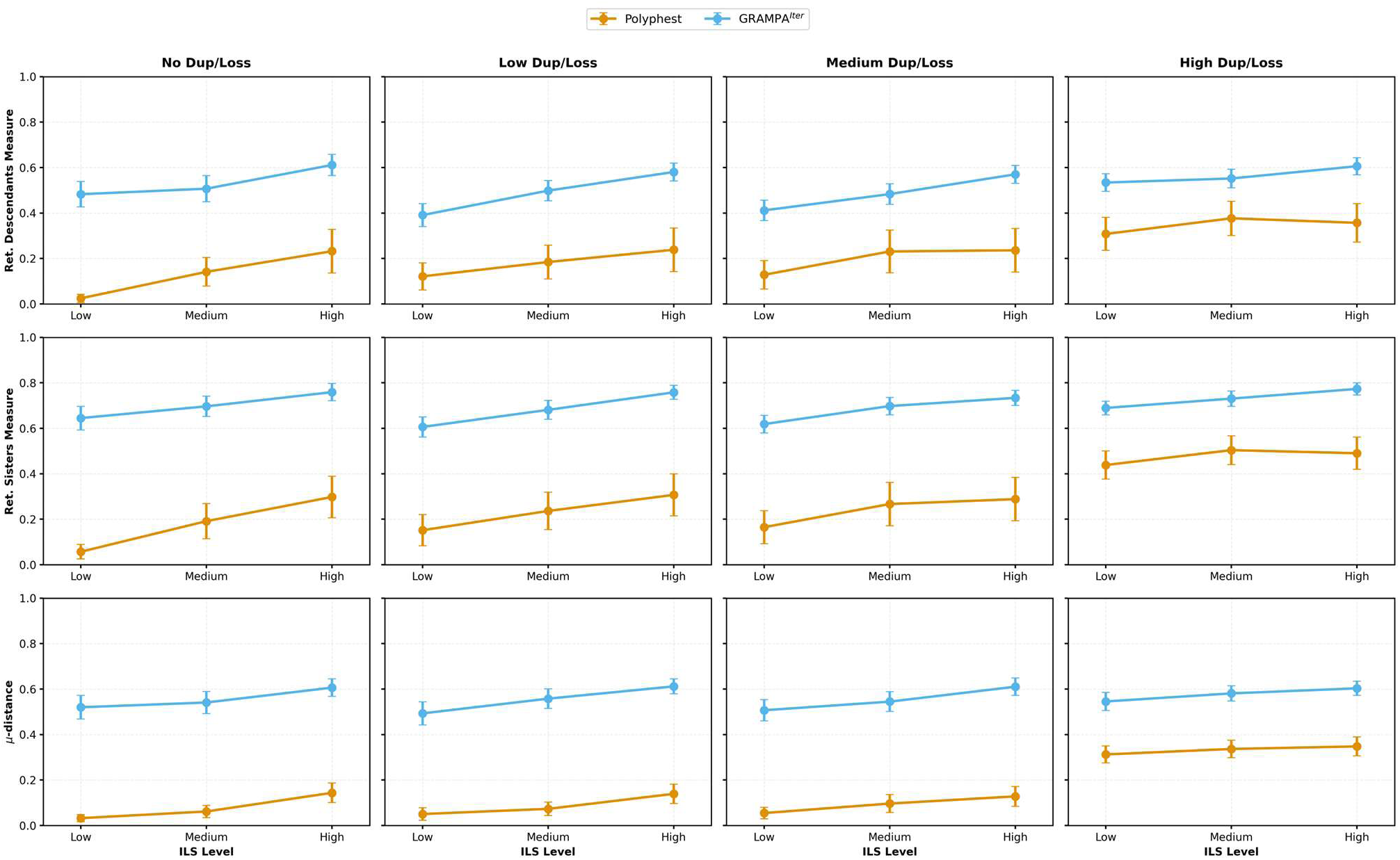
Accuracy of Polyphest and GRAMPA^Iter^ across the 12 baseline configurations. The top row presents the reticulation descendants measure, the middle row the reticulation sisters measure, and the bottom row the μ-distance. Within each row, the four panels represent, from left to right, simulations with no, low, medium, and high gene duplication and loss rates. Within each panel, the x-axis represents low, medium, and high ILS. Orange and blue lines represent Polyphest and GRAMPA^Iter^, respectively. Lower values indicate greater accuracy.

#### Comparison of all methods on the tetraploid subset

*—*To enable broader comparison across methods, we focused on the eight tetraploid networks to which AlloppNET is applicable and on the three ILS configurations without gene turnover, yielding 120 possible simulated datasets per method (Table 4). Among methods with broad completion on this subset, Polyphest again showed the highest accuracy on all three measures. MPAllopp and GRAMPA^Iter^ showed intermediate accuracy and did not differ significantly on any of the three measures across the six networks on which both methods completed (p ≥ 0.44). Within the five-day limit, AlloppNET chains completed on average less than 0.6% of the targeted 10^8^ MCMC iterations, and the resulting networks were among the least accurate. PADRE showed high accuracy among its completed runs, obtaining the lowest reticulation descendants measure and μ-distance. However, these results are based on only 13 simulated datasets and may therefore be biased toward easier inputs, precluding direct comparison with the other methods.

**Table 4.** Accuracy of all methods on the eight tetraploid networks, aggregated across the three ILS configurations in the absence of gene duplication and loss. Values are means across all completed analyses out of 120 simulated datasets (8 networks × 5 replicates × 3 configurations).

| **Method** | **Networks obtained (N/120)** | **Reticulation descendants** | **Reticulation sisters** | **μ-distance** |
|---|---|---|---|---|
| Polyphest | 117 (97.5%) | 0.20 | 0.25 | 0.10 |
| MPAllopp | 88 (73.3%) | 0.33 | 0.65 | 0.44 |
| GRAMPA<sup>Iter</sup> | 119 (99.2%) | 0.48 | 0.61 | 0.37 |
| AlloppNET<sup>a</sup> | 120 (100%) | 0.44 | 0.91 | 0.85 |
| PADRE | 13 (10.8%) | 0.14 | 0.29 | 0.06 |

<sup>a</sup>AlloppNET runs did not converge within the five-day runtime limit

#### The effect of post-WGD fractionation

*—*To more realistically model polyploid genome evolution by incorporating post-WGD gene loss during diploidization, we examined the medium-ILS, medium gene-turnover configuration across four levels of gene retention, from 1.0 (no fractionation) to 0.25 (extensive fractionation). Polyphest’s accuracy in recovering reticulation events was moderately affected at a retention probability of 0.75 but declined sharply with stronger fractionation (Fig. 7). In contrast, GRAMPA^Iter^ was considerably less sensitive to fractionation, showing only a moderate decline in accuracy across the full retention range. Across networks, all three measures increased more steeply with fractionation for Polyphest than for GRAMPA^Iter^ (*p* ≤ 0.01). At 25% retention, Polyphest approached complete disagreement with the true reticulations on both reticulation-specific measures, whereas GRAMPA^Iter^ retained substantially greater accuracy (p < 0.05).

**Figure 7.**
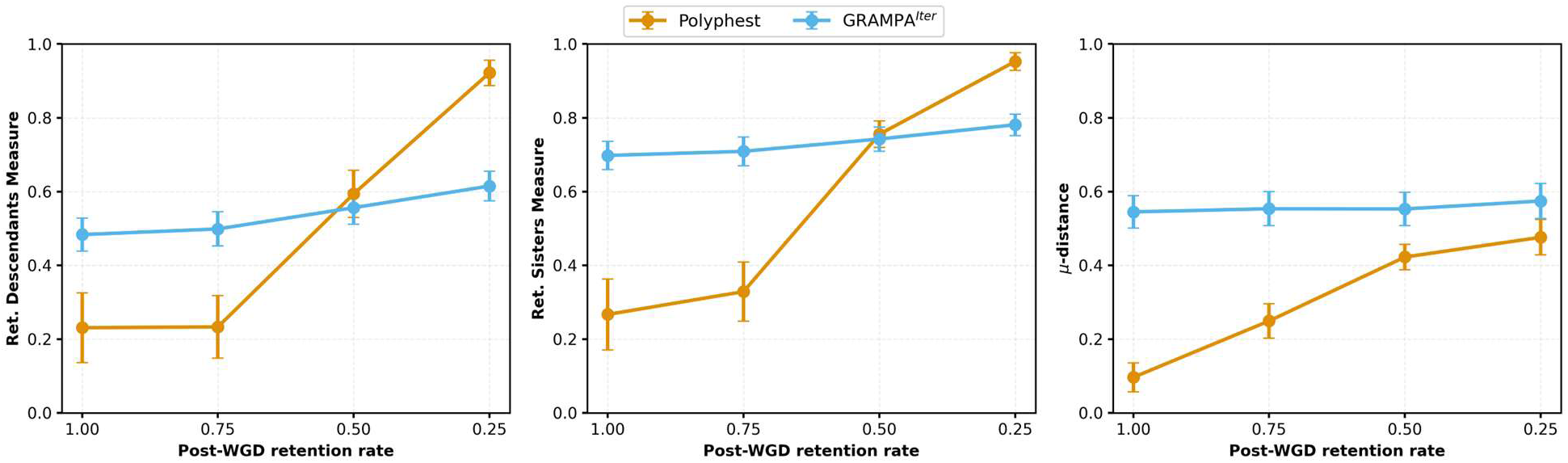
Accuracy of Polyphest and GRAMPA^Iter^ across post-WGD gene retention rates. Fractionation was applied to the medium ILS, medium gene turnover configuration, with the unfractionated configuration shown as a retention rate of 1.00; the x-axis runs from no fractionation on the left to the most severe on the right. (Left) Reticulation descendants measure. (Middle) Reticulation sisters measure. (Right) μ-distance.

The sharp decline in Polyphest’s reticulation accuracy under strong fractionation was compounded by its reliance on inferred ploidy, which became increasingly inaccurate. Ploidy inference correctly identified 100% of polyploid taxa at *q* = 1 and 0.75, but this proportion fell sharply to 40.6% at *q* = 0.5 and to 12.1% at *q* = 0.25. With stronger fractionation, WGD-derived genes were increasingly reduced to single copy, causing ploidy inference from gene-copy number to increasingly classify genuine polyploid species as diploids and exclude them from Polyphest’s search for reticulation events. The effect was also reflected in the number of inferred reticulations, which shifted from overestimation at retention probabilities of 1.0 and 0.75 to substantial underestimation at 0.50 and 0.25. In contrast, GRAMPA^Iter^ does not require ploidy information as input and was therefore not affected by this source of error.

The μ-distance provided a markedly different assessment of method accuracy from the reticulation-specific measures. Although it increased for Polyphest with increasing fractionation, it remained lower than that of GRAMPA^Iter^ across the entire simulated retention range (Fig. 7). At 50% retention, the μ-distance was significantly lower for Polyphest (*p* < 0.05), whereas the reticulation-specific measures did not differ significantly between the methods. At 25% retention, Polyphest had a substantially higher reticulation descendants measure than GRAMPA^Iter^ (0.92 versus 0.61), yet retained a lower μ-distance (0.48 versus 0.57), although this difference was non-significant (*p* = 0.08). Thus, under strong fractionation, the μ-distance did not reflect the reversal in reticulation accuracy revealed by the reticulation-specific measures.

## Discussion

Polyploid phylogenetic inference methods are regularly applied to reconstruct allo- and auto-polyploidization events in large clades with complex evolutionary histories. To date, however, they have been evaluated on small, simulated networks, typically involving only a handful of species and very few polyploidization events (e.g., Jones 2017; Yan et al. 2022, 2024). Their performance on the more complex evolutionary histories found in most empirical studies remains largely unknown. The empirical systems considered here include up to 82 species, multiple polyploidization events, and ploidy levels represented by as many as nine subgenome copies. We compared five existing methods, alongside GRAMPA^Iter^, an automated multi-event extension of GRAMPA developed in this study, across a broad range of empirical polyploid systems and simulations designed to reflect their complexity. Our results reveal substantial differences in accuracy among methods. Polyphest was the most accurate method when gene-copy number reflected ploidy, but it was also more sensitive than GRAMPA^Iter^ to fractionation, a common outcome of genome evolution after polyploidization. On the empirical datasets, the methods often reconstructed markedly different polyploidization histories from the same data.

Previous comparisons of polyploid networks have relied on a single measure of overall topological similarity (e.g., Yan et al. 2022, 2024), which cannot distinguish whether differences between networks arise from spurious or missing reticulation events, the identity of the inferred polyploid lineage, or its parental lineages. The two reticulation-specific measures introduced here separate these components, complemented by the µ-distance as a measure of overall network similarity. This distinction proved informative in practice. Across the empirical datasets, the µ-distance between Polyphest and GRAMPA^Iter^ was lowest for a dataset where the two methods disagreed on almost every reticulation. Similarly, in our simulations under strong fractionation, Polyphest’s µ-distance remained lower than that of GRAMPA^Iter^ despite substantially poorer recovery of the true polyploidization events. Because the µ-distance summarizes the entire network, disagreement in the largely non-reticulate portion of the network contributes similarly to disagreement in the polyploidization events themselves. Our results thus demonstrated that overall network similarity is not a reliable proxy for agreement or accuracy in the inferred reticulation events.

Across the baseline simulations, both ILS and gene turnover reduced inference accuracy, although gene turnover generally had the stronger effect. Polyphest was particularly sensitive to increasing gene turnover, whereas GRAMPA^Iter^ showed a more moderate response but remained less accurate across all baseline configurations. GRAMPA^Iter^’s lower accuracy may partly reflect limitations of its inference framework. Apart from its reliance on a correct input species tree, GRAMPA^Iter^ cannot detect some types of nested events. The divide- and-conquer approach introduced here assumes independence between the sub-problems created after each inference event. However, when two sequential events are nested with respect to only one of their parental lineages, the order in which the events are inferred may prevent detection of the second event.

The fractionation simulations capture an important characteristic of polyploid genome evolution that is absent from the baseline simulations: during diploidization, extensive loss of WGD-derived gene copies progressively erodes the correspondence between gene-copy number and the ploidy level (Van de Peer et al. 2017; Li et al. 2021). Our simulations approximate the outcome of this process by instantaneously removing a defined fraction of WGD-derived copies. Under this scenario, the decline in Polyphest’s accuracy was compounded by its reliance on prior ploidy information. When ploidy is inferred from gene-copy numbers, polyploids that have lost most of their duplicated gene copies may be classified as diploids and excluded from Polyphest’s search for reticulation events, whereas GRAMPA^Iter^ does not require ploidy as input and is not exposed to this source of error. More generally, any method that identifies polyploids from gene-copy number inherits the loss of this signal during diploidization, so its accuracy is expected to decrease with the age of the polyploidization event and as diploidization progresses.

Ploidy information obtained independently of gene trees, for example through flow cytometry, chromosome counts, or sequencing reads, may be less sensitive to this source of error (Doležel et al. 2007; Weiß et al. 2018; Gaynor et al. 2024; Shafir et al. 2025). Whole-genome sequencing can provide additional information, since patterns of variation among chromosome copies can help distinguish autotetraploids from allotetraploids (Ranallo-Benavidez et al. 2020). Such information could be more informative than ploidy alone, yet it is currently not utilized by any existing method. Like gene-copy number, however, these sources are most informative for recent WGD events, since DNA loss and chromosomal rearrangements following polyploidization gradually obscure features of the original polyploid genome (Leitch and Bennett 2004; Mandáková and Lysak 2018; Li et al. 2021). Moreover, even accurate ploidy information did not resolve the disagreement among methods. In the *Ephedra* dataset (Wu et al. 2016) for example, Polyphest and AlloppNET were constrained by the same flow-cytometry-based ploidy assignments but still inferred substantially different reticulation histories. Ploidy information can therefore constrain which taxa are considered polyploid, but not their inferred evolutionary origins.

Resolving polyploid phylogenetic placement may require integrating signals beyond gene trees. Distributions of synonymous divergence (Ks) between paralogs can identify WGDs and position them relative to speciation events (Zwaenepoel and Van de Peer 2019; Sensalari et al. 2022), while genomic collinearity, gene content, and microsynteny can identify WGD-derived copies and distinguish subgenomes (Emery et al. 2018; Sun et al. 2022; Ding et al. 2023). These signals complement that obtained by gene trees, as Ks distributions and synteny provide evidence for the occurrence and timing of WGD, whereas gene trees can identify potential progenitor lineages but rely on duplicated copies that are progressively lost. Integrating these sources with independent ploidy estimates could therefore constrain inference where any single source is weak. Machine learning could offer a natural framework for such integration, since training data with known histories can be generated in large quantities, and neural networks trained on coalescent simulations have already been used to select among hybridization scenarios (Blischak et al. 2021). The accuracy of such an approach would depend on training simulations that capture the complexity of empirical data (Braichenko et al. 2025), and the simulation framework developed here could provide a basis for generating such training data.

Several limitations should be considered when interpreting these results. Not all methods could be evaluated equally. All methods were run under the same five-day limit per analysis, within which AlloppNET did not converge and PADRE and MPAllopp completed only a small and potentially biased subset of simulations. In addition, five model networks used for simulation were inferred by methods included in the benchmark. Although gene trees were subsequently simulated independently, these model trees may retain method-specific characteristics, such as differences in whether reticulations tend to be placed toward the tips or deeper in the phylogeny. Our fractionation model, based on the retention framework of Rabier et al. (2014), provides a simplified representation of post-WGD gene loss by applying a fixed retention probability to WGD-derived copies. Moreover, all simulations used 1,000 gene trees and relied on the taxon sampling of the model networks, so we did not assess how accuracy depends on these data characteristics. Incorporating additional biological complexity into the simulations would likely make the inference problem more challenging, and thus the accuracies reported here may represent optimistic estimates of performance on complex empirical histories.

Taken together, our results demonstrate substantial uncertainty in the reconstruction of complex polyploid histories and show that method performance depends strongly on the evolutionary processes shaping the data. The comparison framework introduced here provides a means to identify the sources of disagreement among inferred networks and should facilitate more informative evaluation of polyploid phylogenetic inference methods.

## Supplementary Material

Supplementary Methods, Supplementary Tables S1–S2 and Supplementary Figures S1–S5 are available in the Supplementary Material file accompanying this preprint.

## Acknowledgments

We thank Z. Yan and L. Nakhleh for advice on running Polyphest and MPAllopp.

## Data Availability

The code for the simulations, the method adapters, the evaluation and the figures and tables of this study is available at https://github.com/tomulanovski/polyploid-benchmark. The simulated and empirical datasets and the inferred networks will be deposited in the Dryad Digital Repository upon publication.

