## Supplementary Material for "How Reliable Are Reconstructions of Complex Polyploid Histories? A Benchmark of Phylogenetic Inference Methods"

Contents: Supplementary Methods; Supplementary Tables S1–S2; Supplementary Figures S1–S5; Supplementary References.

#### Supplementary Methods

Detailed descriptions of the five polyploid inference methods summarized in the main text (Methods Evaluated) and in Table 1.

##### PADRE

The method receives as input a collection of gene trees that are represented as multi-labeled trees (MUL-trees), defined as phylogenetic trees in which some species appear multiple times. This structure is well-suited for polyploid taxa because it reflects the multiple gene copies present in genomes of varying ploidy: diploid species typically have one gene copy, tetraploids have two copies, hexaploids have three copies, etc. Each copy corresponds to a distinct subgenome that can be traced to different ancestral lineages.

PADRE employs a two-step procedure to infer a phylogenetic network from a collection of MUL-trees. The first step involves the construction of a consensus MUL-tree from the input gene trees. In this step, clusters (i.e., sets of leaf labels that are descended from a specific branch) are extracted from the input MUL-trees. The clusters are classified as core clusters and ambiguous clusters (Supplementary Fig. S3): core clusters contain at least one taxon that appears only once across the gene trees, so when such a cluster can be added to the consensus MUL-tree under construction, there is only one position in which it can be inserted. Ambiguous clusters, in contrast, consist solely of multiply labeled taxa and can usually be inserted into the consensus MUL-tree in several different positions, each yielding a different MUL-tree. Core and ambiguous clusters are ranked separately according to their level of support (i.e., the number of input MUL-trees in which they occur) and considered sequentially in a two-phase greedy fashion to construct a consensus MUL-tree. According to these rankings, core clusters are first considered followed by consideration of the ambiguous clusters. In each iteration, the considered cluster is added only if it is compatible with the current tree. This process continues until no further clusters can be added without violating compatibility. The result of this step is a consensus MUL-tree.

The second step involves the construction of a phylogenetic network from the consensus MUL-tree. PADRE aims to construct a phylogenetic network that explains the MUL-tree with the minimum number of hybridization events. Specifically, it iteratively identifies a collection of equivalent subtrees, meaning subtrees with identical topologies and leaf labels, within the consensus MUL-tree. Each such collection of subtrees is then merged, such that redundant copies of the subtrees are pruned while preserving their connections to the original ancestral lineages. This operation introduces a reticulation node, having in-degree of more than one, thus representing a hybridization event. Two examples of this merging operation are shown in Supplementary Fig. S4. This is done iteratively until no equivalent subtrees remain.

#### AlloppNET

AlloppNET (Jones et al. 2013; Jones 2017) is a Bayesian method designed for inferring species MUL-trees in the presence of both polyploidy and ILS. AlloppNET operates directly on sequence alignments. It additionally requires an input file that assigns each sequence to a species and indicates whether that species is diploid or tetraploid. When the homeologous relationships among an allotetraploid's sequences are known, this information can also be supplied, and in this case AlloppNET determines the optimal homeolog assignment during its search.

AlloppNET uses a reversible-jump Markov Chain Monte Carlo (MCMC) algorithm with MCMC operations that can swap sequence assignments between parental species and explore different topologies. The main restriction of AlloppNET is that it only considers diploid and allotetraploid species (ploidy levels 2 and 4). This limitation means that clades containing species with higher ploidy levels, e.g., hexaploids and octoploids, cannot be analyzed.

#### GRAMPA

GRAMPA (Gene-tree Reconciliation Algorithm for MUL-trees and Polyploid Analysis) (Thomas et al. 2017) represents a different approach for polyploid phylogeny reconstruction based on the well-established last common ancestor (LCA) reconciliation principle (Goodman et al. 1979; Page 1994). In its classic formulation, LCA mapping reconciles a gene tree with a singly labeled species tree by inferring the minimum number of gene duplication and loss events required to explain the gene tree. GRAMPA extends this framework by allowing LCA mapping to be performed for MUL-trees with a single duplicated lineage. This addresses a fundamental challenge because, in MUL-trees, some nodes have multiple possible mappings since tips in the species tree are not uniquely labeled. GRAMPA solves this by evaluating all possible mappings of genes from polyploid species and selecting the mapping that minimizes the reconciliation score, defined as the sum of inferred gene duplication and loss events across all gene trees.

The core functionality involves identifying the optimal placement of a polyploidy event on a phylogeny. To this end, GRAMPA defines two key nodes in the species tree that together specify a candidate polyploidization scenario: H1 marks the location of the polyploid clade in the singly-labeled species tree, and H2 marks the branch onto which a copy of the H1 subtree is grafted to construct the MUL-tree (Supplementary Fig. S5). After grafting, each sub-genome copy is placed as a sister clade to one of the two parental lineages: the clade originally sister to H1 is the first parental lineage, and the clade subtended by H2 is the second. By testing different combinations of H1 and H2 placements, GRAMPA generates multiple candidate MUL-trees and identifies which scenario yields the lowest reconciliation score. Notably, the placement of H1 and H2 in the optimal MUL-tree reveals both the type of polyploidy and its phylogenetic context. Placement of H1 and H2 on different lineages indicates allopolyploidy, whereas placement on the same node is indicative of autopolyploidy.

A major limitation of the current implementation of GRAMPA is that it can infer only a single polyploidization event or none if the reconciliation score of the input species tree is lower than all alternative MUL-trees. This is because each candidate MUL-tree is constructed by a single graft of the H1 subtree onto the branch subtended by H2, which results in exactly one WGD. Consequently, the method is not designed to detect multiple or sequential polyploidy events and cannot be used for complex systems involving species with ploidy levels higher than tetraploidy. As a result, the method has limited applicability to clades with complex evolutionary histories involving repeated polyploidization. An additional limitation is that the reconciliation score weights gene duplications and losses equally, which may be problematic in polyploid lineages where diploidization typically drives extensive gene loss exceeding gains.

#### MPAllopp

MPAllopp (Maximum Parsimony for Allopolyploids) (Yan et al. 2022) is also based on the reconciliation principle but it uses a different parsimony approach than GRAMPA for inferring phylogenetic networks. MPAllopp extends the Minimizing Deep Coalescences (MDC) criterion to account for polyploidy while simultaneously handling ILS (Maddison 1997). The MDC approach seeks phylogenetic hypotheses that minimize the number of these deep coalescence events, thereby providing the most parsimonious explanation for observed gene tree incongruence.

The most recent common ancestor (MRCA) of two homeologs predates the hybridization event and resides in two different parental lineages. Thus, the two genes are only expected to coalesce at the deeper ancestors where their respective subgenomes meet, at or above the hybridization node. MPAllopp recognizes that gene copies from different subgenomes are not expected to coalesce within the polyploid species itself, since they originated from separate parental ancestors. Deep coalescences are therefore counted only where the coalescent model genuinely predicts them, and not as an artifact of polyploidy. MPAllopp computes its parsimony score by summing the extra lineages across all branches of the MUL species tree. For any given branch, if $k$ lineages enter from the descendants but do not merge, the score for that branch is $k - 1$. A score of zero represents a perfect fit where all lineages coalesce as expected.

MPAllopp employs a heuristic search strategy to explore the space of possible networks, applying various topological modifications and evaluating each candidate based on its parsimony score. Due to the heuristic nature of its search algorithm, MPAllopp is not guaranteed to find the optimal network in a single run. Therefore, it is recommended to execute the method multiple times and select the best solution among all runs. This approach helps mitigate the risk of getting trapped in local optima during the network space exploration.

#### Polyphest

The PADRE algorithm described above has a notable limitation: it does not account for ILS, an evolutionary process that can generate discordance among gene trees even in the absence of hybridization events. Polyphest (POLYploid PHylogeny ESTimation) extends PADRE's algorithm to jointly accommodate ILS and polyploidy while maintaining computational efficiency (Yan et al. 2024). Polyphest follows the same two-step structure as PADRE, with modifications to both steps designed to make the inference robust to ILS.

To reduce the computational burden of cluster selection, which becomes NP-hard for MUL-trees, Polyphest introduces the concept of "approximate compatibility" between clusters as a relaxation of strict compatibility. The strict rule, which works for uniquely labeled trees, considers two clusters compatible only if one is nested within the other or if they are disjoint as sets. This rule produces false rejections on MUL-trees, because it treats labels as identifiers of unique leaves and ignores the fact that the same label can appear on multiple leaves. For example, the clusters {x, y} and {x, z} are neither nested nor disjoint, so the strict rule rejects them, even though both can coexist in a MUL-tree in which the two copies of x represent its two different subgenome copies. Polyphest's relaxed criterion handles this case directly: two clusters are approximately compatible if one cluster is nested within the other, or if they do not compete for the same copies of taxa. The latter condition replaces strict set-disjointness with a weaker requirement that still respects the multiplicity of labels in the target MUL-tree.

In the first step of reconstructing the consensus MUL-tree, Polyphest ranks and selects the optimal set of clusters by solving the Maximum Weight Approximate Compatible Clusters problem using integer linear programming. This process identifies the best collection of approximately compatible clusters, weighted by factors including cluster multiplicity, frequency across gene trees, and cluster size. The selected group of clusters is then used to build a consensus MUL-tree. Unlike PADRE, Polyphest does not prioritize core clusters over ambiguous clusters, based on the observation that ILS can weaken the reliability of core clusters as indicators of true evolutionary relationships, and that including core clusters could block the inclusion of other informative, highly supported ambiguous clusters.

In the second step, network construction largely follows that of PADRE network building algorithm, with one main modification to accommodate ILS. Instead of requiring equivalent subtrees to be strictly identical in topology and leaf labels, Polyphest allows them to be nearly identical, controlled by a user-defined threshold $\delta$. Specifically, two subtrees are treated as equivalent when their normalized graph edit distance does not exceed $\delta$. Higher values of $\delta$ permit more topological variation between merged subtrees. This allows slightly different subtrees to be considered as representing the same polyploidization event, thereby accounting for topological variations that may arise due to ILS. When the threshold is set to zero, Polyphest reduces to PADRE's network construction algorithm.

Aside from algorithmic differences to PADRE, Polyphest requires as an additional input the specification of the leaf labels in the target MUL-tree. This means that the user should specify the number of copies of each taxon in the consensus MUL-tree and thus must make prior assumptions regarding the ploidy levels of the species being analyzed. For example, the user must declare in advance that a hexaploid taxon should appear three times in the reconstructed MUL-tree, once per subgenome, while a diploid taxon should appear once.

### Supplementary Tables

***Supplementary Table S1**. Summary of the 21 empirical datasets.*

| Study | Citation | No. Gene Trees | In 12-dataset comparison | AlloppNET applicable |
|---|---|---|---|---|
| Bendiksby_2011 | (Bendiksby et al. 2011) | 1 | No | No |
| Koenen_2021 | (Koenen et al. 2021) | 4,371 | Yes | Yes |
| Brysting_2007 | (Brysting et al. 2007) | 1 | No | No |
| Lawrence_2016 | (Lawrence and Datwyler 2016) | 2 | Yes | No |
| Diaz-Perez_2018 | (Díaz-Pérez et al. 2018) | 3 | Yes | No |
| Wisecaver_2023 | (Wisecaver et al. 2023) | 2,699 | Yes | No |
| Ding_2023 | (Ding et al. 2023) | 4,653 | Yes | Yes |
| Liang_2021 | (Liang et al. 2021) | 1 | No | No |
| Popp_2005 | (Popp et al. 2005) | 6 | Yes | No |
| Wu_2016 | (Wu et al. 2016) | 2 | Yes | Yes |
| Liu_2023 | (Liu et al. 2023) | 1 | No | No |
| Ren_2024 | (Ren et al. 2024) | 727 | Yes | No |
| Marcussen_2011 | (Marcussen et al. 2011) | 1 | No | No |
| Marcussen_2012 | (Marcussen et al. 2012) | 4 | Yes | No |
| Sessa_2012 | (Sessa et al. 2012) | 1 | No | No |
| Zhao_2021 | (Zhao et al. 2021) | 982 | Yes | Yes |
| Hori_2014 | (Hori et al. 2014) | 1 | No | No |
| Marcussen_2015 | (Marcussen et al. 2015) | 3 | Yes | No |
| Shahrestani_2015 | (Mohammadi Shahrestani et al. 2015) | 1 | No | No |
| Morales-Briones_2021 | (Morales-Briones et al. 2021) | 923 | Yes | No |
| Soza_2014 | (Soza et al. 2014) | 1 | No | No |

***Supplementary Table S2****. Summary of the 21 model networks used in simulations. Max. copies is the largest number of subgenome copies of any taxon; No. Hybridization events and No. Autopolyploid events sum to Total WGDs.*

| Study | No. Taxa | No. Polyploids | Max. Copies | No. Hybridization events | No. Autopolyploid events | Total WGDs | Network source |
|---|---|---|---|---|---|---|---|
| Bendiksby_2011 | 40 | 6 | 2 | 6 | 0 | 6 | Original study |
| Koenen_2021 | 18 | 5 | 2 | 1 | 0 | 1 | GRAMPA |
| Brysting_2007 | 27 | 17 | 3 | 19 | 0 | 19 | IQ-TREE |
| Lawrence_2016 | 38 | 5 | 3 | 5 | 0 | 5 | Original study |
| Diaz-Perez_2018 | 18 | 9 | 3 | 10 | 3 | 13 | Polyphest |
| Wisecaver_2023 | 15 | 1 | 2 | 1 | 0 | 1 | Original study |
| Ding_2023 | 12 | 7 | 2 | 0 | 1 | 1 | Original study |
| Liang_2021 | 31 | 24 | 4 | 29 | 0 | 29 | Original study |
| Popp_2005 | 13 | 3 | 3 | 2 | 2 | 4 | PADRE |
| Wu_2016 | 18 | 9 | 2 | 8 | 0 | 8 | PADRE |
| Liu_2023 | 26 | 8 | 2 | 6 | 2 | 8 | Original study |
| Ren_2024 | 16 | 7 | 4 | 5 | 0 | 5 | Original study |
| Marcussen_2011 | 20 | 7 | 5 | 10 | 4 | 14 | Original study |
| Marcussen_2012 | 39 | 35 | 9 | 11 | 0 | 11 | Original study |
| Sessa_2012 | 54 | 16 | 3 | 24 | 0 | 24 | IQ-TREE |
| Zhao_2021 | 16 | 14 | 2 | 1 | 0 | 1 | Original study |
| Hori_2014 | 16 | 7 | 3 | 11 | 0 | 11 | IQ-TREE |
| Marcussen_2015 | 16 | 13 | 6 | 21 | 0 | 21 | Original study |
| Shahrestani_2015 | 5 | 2 | 2 | 2 | 0 | 2 | IQ-TREE |
| Morales-Briones_2021 | 82 | 49 | 3 | 49 | 0 | 49 | Polyphest |
| Soza_2014 | 52 | 22 | 9 | 35 | 1 | 36 | Original study |

#### Supplementary Figures

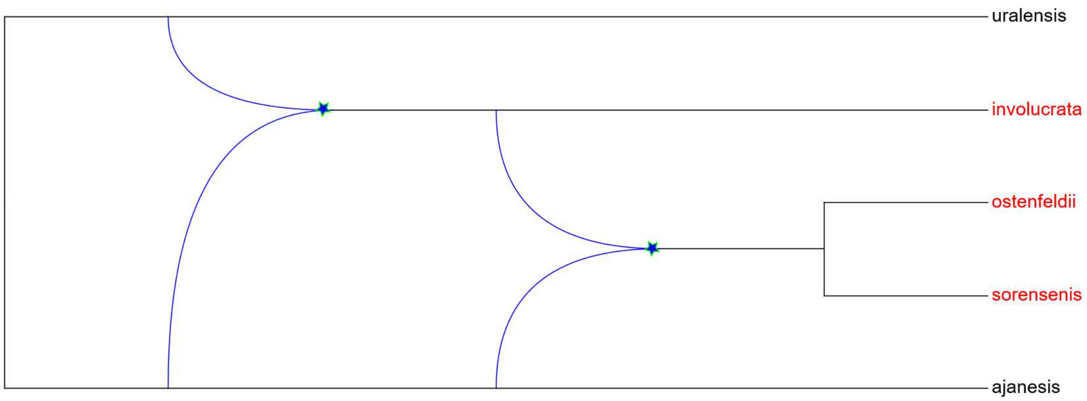


***Supplementary Figure S1***. *Phylogenetic network for the Silene dataset as inferred by Popp et al. (2005).*

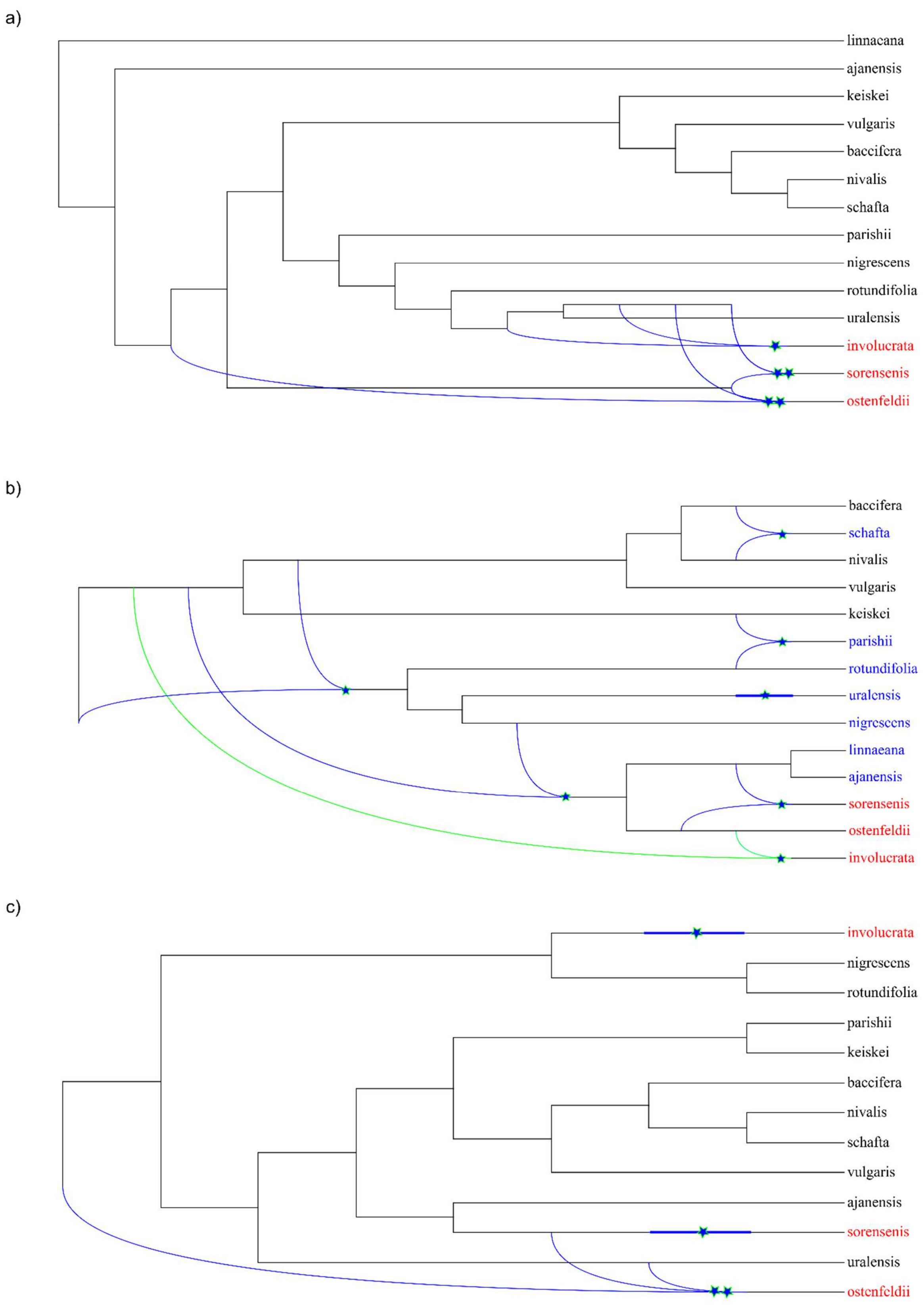
a)
linnaeana
ajanensis
keiskei
vulgaris
baccifera
nivalis
schafta
parishii
nigrescens
rotundifolia
uralensis
involucrata
sorensenis
ostenfeldii
b)
baccifera
schafta
nivalis
vulgaris
keiskei
parishii
rotundifolia
uralensis
nigrescens
linnaeana
ajanensis
sorensenis
ostenfeldii
involucrata
c)
involucrata
nigrescens
rotundifolia
parishii
keiskei
baccifera
nivalis
schafta
vulgaris
ajanensis
sorensenis
uralensis
ostenfeldii

***Supplementary Figure S2*. *Inferred phylogenetic networks for the Silene dataset from Popp et al. (2005).*** *(a) Polyphest (5 reticulation events, 3 polyploids). (b) GRAMPA$^{Iter}$ (7 events, 10 polyploids); the green reticulation denotes the event inferred in its first iteration. (c) PADRE (4 events, 3 polyploids). Stars denote branches with whole-genome duplication events (autopolyploidization or allopolyploidization). Red taxa indicate the three polyploid species (S. involucrata, S. sorensenis, S. ostenfeldii) identified in the original study and provided as prior information to Polyphest. Blue taxa indicate additional species inferred as polyploid exclusively by GRAMPA$^{Iter}$.*

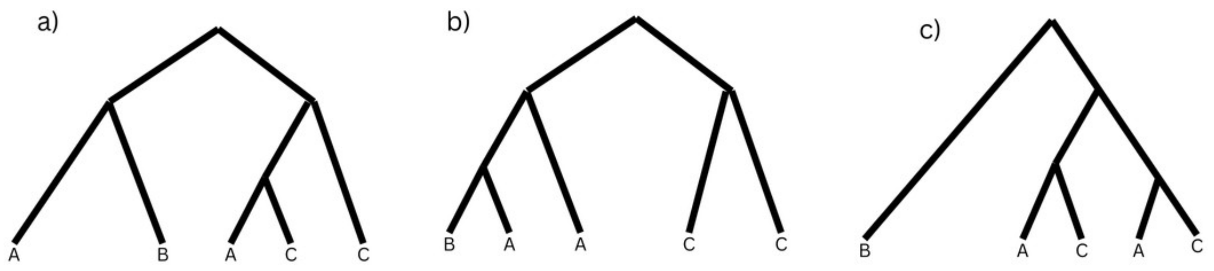


***Supplementary Figure S3. Core and ambiguous clusters.*** *Three gene trees represented as MUL-trees, in which some taxa appear more than once. The set of all clusters obtained from these trees is partitioned into core and ambiguous clusters. Core clusters contain at least one uniquely labeled taxon and can be inserted into the consensus MUL-tree in a unique way, whereas ambiguous clusters consist solely of multiply labeled taxa and may have multiple placements. Across the three tree inputs, B is the only uniquely labeled taxon and the core clusters are {A,A,B} and {A,B}, whereas the ambiguous clusters are {C,C}, {A,C,C}, {A,C}, and {A,A,C,C}.*

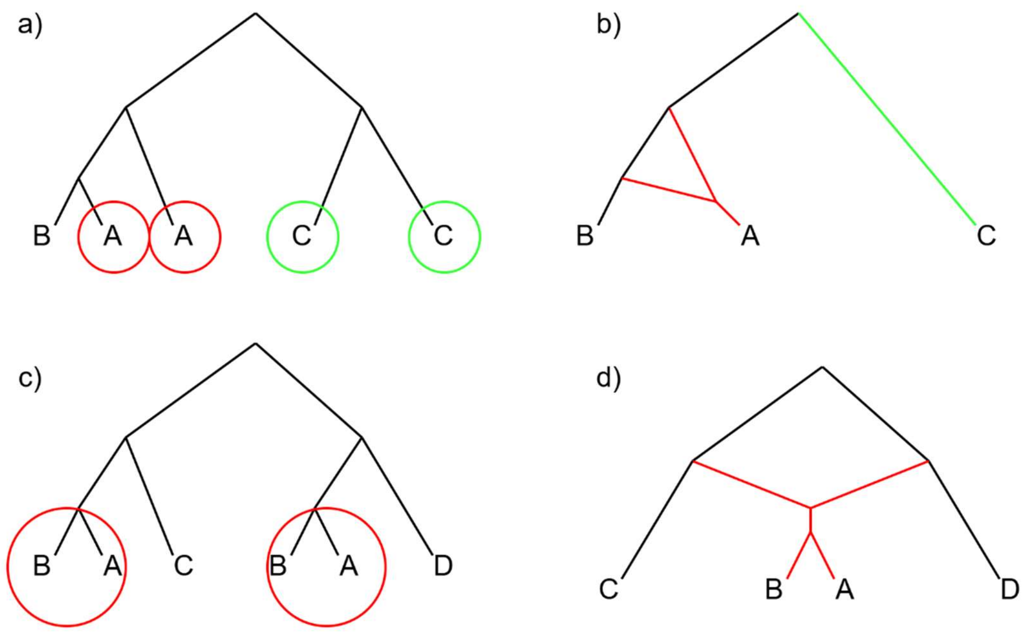

***Supplementary Figure S4*. *Construction of phylogenetic networks from consensus MUL-trees using PADRE.*** *The figure illustrates two examples of converting MUL-trees into networks. In each case, the circled subtrees are the equivalent subtrees that are merged, and merging prunes the redundant copies while retaining their connections to the original ancestral lineages, introducing a reticulation that represents the inferred hybridization event.* ***a)*** *The consensus MUL-tree obtained from the collection of MUL-trees in Supplementary Fig. S3.* ***b)*** *The resulting network. The red edges represent an allopolyploid event resulting from the hybridization of lineages A and B, while the green edges represent an autopolyploid event within lineage C.* ***c)*** *A second consensus MUL-tree.* ***d)*** *The resulting network after merging.*

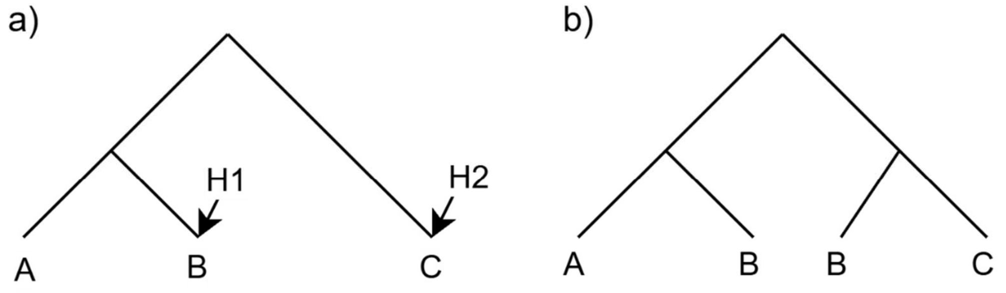


***Supplementary Figure S5*. *Placement of H1 and H2 nodes***. *a) The H1 node in the singly labeled species tree indicates that species B may be the result of a polyploidization event. The H2 node in the singly labeled species tree indicates the second parental lineage of the polyploid species B. b) The MUL species tree incorporating these H1 and H2 hypotheses.*
